# Antisense oligonucleotide allele-specific targeting of EFEMP1 in a patient-derived model of Doyne honeycomb retinal dystrophy

**DOI:** 10.1101/2025.07.16.664883

**Authors:** Farah. O. Rezek, Beatriz Sanchez-Pintado, Emily R. Eden, Nancy Aychoua, Andrew R. Webster, Amanda-Jayne F. Carr, Michel Michaelides, Michael E. Cheetham, Jacqueline van der Spuy

## Abstract

Doyne honeycomb retinal dystrophy is an incurable juvenile macular dystrophy that leads to visual impairment by early to mid-adulthood. It is an autosomal dominant disorder caused by a c.1033C>T, p.Arg(345Trp) variant in *EFEMP1*, and is characterised by the early onset extracellular deposition of drusen between the retinal pigment epithelium basement membrane and underlying layers of Bruch’s membrane. In this study, we developed an antisense oligonucleotide approach to target *EFEMP1*. We reprogrammed patient-derived renal epithelial cells to induced pluripotent stem cells followed by directed differentiation to retinal pigment epithelium and compared the phenotype to gene-corrected and *EFEMP1* knockout patient-derived retinal pigment epithelium. In the patient-derived disease model, remodelling of the extracellular matrix occurred with progressive accumulation of extracellular deposits containing the drusen-associated proteins apolipoprotein E and collagen IV, in addition to EFEMP1. Moreover, the intracellular accumulation of neutral lipids was evident. We developed an allele-specific antisense oligonucleotide which specifically and effectively promoted the clearance of the *EFEMP1* c.1033C>T transcript in the patient-derived disease model following assisted or gymnotic delivery. In this disease model, gymnotic delivery led to a decrease in extracellular deposits and cleared the intracellular accumulation of lipids, even after the onset of this disease phenotype, suggesting this could be a practical and effective therapeutic approach.

## INTRODUCTION

Doyne honeycomb retinal dystrophy (DHRD) is a progressive and incurable juvenile macular dystrophy that is characterised by the early onset radial deposition of peripapillary and macular drusen that merge over time in a ‘honeycomb’ pattern.^1,2^ The lipoproteinaceous drusen are deposited extracellularly between the retinal pigment epithelium (RPE) basement membrane and underlying layers of Bruch’s membrane. Though the onset of visual symptoms varies, DHRD typically leads to visual impairment in early to mid-adulthood. However, extensive drusen can be present as early as the second decade of life.^1,2^ Early visual symptoms include decreased visual acuity, photophobia, metamorphopsia, dyschromatopsia and relative scotomas that progress to atrophy of the retinal pigment epithelium (RPE) leading to central visual loss. A worse prognosis is associated with choroidal neovascularisation (CNV).^3^

DHRD is an autosomal dominant disorder caused by the c.1033C>T, p.(Arg345Trp) variant in the *epidermal growth factor (EGF)-containing fibulin-like extracellular matrix protein 1* (*EFEMP1*) gene.^4^ The EFEMP1, or fibulin 3 (F3), protein is an extracellular glycoprotein that is synthesised by the RPE and secreted into the collagen IV-rich RPE basement membrane where it contributes to the structural integrity of the extracellular matrix (ECM). The Arg345Trp substitution is in the final Ca^2+^ binding EGF domain of EFEMP1, which alters disulphide bond formation and the structural integrity of the protein, resulting in increased stability and accumulation of the mutant protein in the ECM.^5–8^ Extracellular accumulation of EFEMP1 Arg345Trp also causes increased expression of its binding partner, tissue inhibitor of metalloproteinase 3 (TIMP3), resulting in reduced matrix metalloproteinases (MMP) activity, such as MMP2 and MMP9, and creating an imbalance that results in the accumulation of ECM aggregates.^9,10^ The remodelling process triggers an interplay that increases levels of complement component 3 (C3) and complement factor B (CFB) that exacerbates the ECM remodelling and sub-RPE deposition.^11–13^ Moreover, EFEMP1 Arg345Trp has been shown to exert a hyper-inhibitory effect on epidermal growth factor receptor (EGFR) signalling, potentially through the deactivation of the PI3K/AKT pathway. This disruption leads to reduced expression of carboxylesterase 1 (CES1), impairing cholesterol efflux and resulting in abnormal lipid accumulation within RPE cells and the basement membrane.^14^ These pathological changes collectively disrupt the structural and functional integrity of the RPE basement membrane leading to the deposition of drusen containing components such as apolipoprotein E (APOE), C3 and TIMP3. Increased accumulation of sub-RPE basal deposits rich in lipids and proteins, including APOE, TIMP3 and EFEMP1, has been reported in a patient-derived DHRD RPE disease model.^13^

Patients that harbour biallelic loss-of-function *EFEMP1* variants develop a pronounced connective tissue disease characterised by multiple and recurrent abdominal and thoracic herniae, myopia, hypermobile joints, scoliosis, and thin translucent skin, however macular dystrophy is absent in these patients.^15^ Moreover, macular dystrophy is absent in *Efemp1* knockout mice and the knockout of *Efemp1* has been shown to be protective against the development of sub-RPE deposits.^16,17^ These data support an EFEMP1 Arg345Trp toxic gain-of-function and the absence of haploinsufficiency in DHRD.

Therefore, in this study, we designed an allele-specific antisense oligonucleotide (ASO) therapy to induce the knockdown of variant *EFEMP1* expression in a patient-derived disease model of DHRD. We reprogrammed patient renal epithelial cells to induced pluripotent stem cells (iPSCs) followed by directed differentiation to RPE (iRPE). Our patient iRPE model faithfully recapitulated the disease phenotype, including the accumulation of lipids, remodelling of the ECM and the sub-RPE deposition of drusen-associated components. The treatment of patient-derived iRPE with therapeutic ASO resulted in allele-specific targeting of *EFEMP1* Arg345Trp and efficient resolution of pathogenic changes, even after the onset of the disease phenotype *in vitro*.

## RESULTS

### DHRD patient clinical phenotype

Renal epithelial cells were purified from urine of a 60-year-old male patient clinically diagnosed with DHRD and confirmed to harbour the autosomal dominant *EFEMP1* variant c.1033C>T, p.Arg(345Trp). The patient was first diagnosed with DHRD in his early twenties, presymptomatically, following retinal examination due to the family history. Ultra-widefield fundus images (Optos scanning laser ophthalmoscope) and autofluorescence images showed hyperautoflourescent deposits at the macula, some with a radial configuration, with deposit directly adjacent to the optic disc (Figure S1). Heidelberg Spectralis optical coherence tomography (OCT) images showed extensive, hyperreflective deposit underlying the neurosensory retina, greater at the left macula than the right (Figure S1).

### Correction of *EFEMP1* c.1033C>T and *EFEMP1* knockout in patient iPSC

The DHRD patient renal epithelial cells were reprogrammed to induced pluripotent stem cells (iPSCs) as described previously.^18,19^ CRISPR-Cas9 homology directed repair (HDR) using *in vitro* assembled *EFEMP1* crRNA/tracrRNA-Cas9 ribonucleotide protein (RNP) together with a single stranded oligonucleotide (ssODN) repair template was used to correct the c.1033C>T, p.Arg(345Trp) variant in the patient iPSC line thereby generating an isogenic control (Figure S2A). In addition, the ssODN repair template introduced a heterozygous synonymous change in the protospacer adjacent motif (PAM). CRIPSR-Cas9 non homologous end joining (NHEJ) targeting *EFEMP1* exon 5 introduced a homozygous insertion of a single nucleotide (G) inducing a frame shift and premature translation termination of the *EFEMP1* transcript (c.215_216insG, p.Lys73GlufsX81) (Figure S2B). crRNA and ssODN sequences are listed in Supplemental Tables 1 and 2 (Table S1, Table S2). No off-target editing was observed in the isogenic lines at the top 10 predicted off-target genomic loci (Table S3). iPSC cultures from the DHRD patient iPSC (R345W), isogenic repaired iPSC (ISOCON) and isogenic knockout (ISOKO) lines uniformly expressed pluripotency markers (*OCT4*, *NANOG*, *TRA1-181*, and *SSEA4*) (Figure S2C).

### Phenotypic characteristics of patient iPSC-iRPE (iRPE)

iPSC lines were differentiated to RPE by the method described in Regent *et al.* (2019).^20^ Light microscopy images taken of the isogenic control (ISOCON) and R345W patient line at day 60 of RPE differentiation (Figure 1A), demonstrated a typical RPE cobblestone morphology, with pronounced cell borders. The ISOCON iRPE formed a homogenous monolayer, while the R345W iRPE displayed a more heterogenous morphology. Immunofluorescence (IF) analysis of both iRPE lines demonstrated clear expression of the tight junction protein ZO-1, which localized to the cell-cell borders in a continuous, hexagonal pattern consistent with the formation of an epithelial monolayer with tight junctions (Figure 1A). PMEL, a marker of melanosome biogenesis, was also detected in both lines, appearing as intracellular puncta indicative of developing pigment organelles (Figure 1A). These data confirm the expression of key RPE markers associated with epithelial identity and early melanosome formation. Western blot analyses of both lines demonstrated robust expression of the RPE characteristic markers ZO-1, Ezrin and MERTK at day 110 of differentiation (Figure 1B).

**Figure 1.**
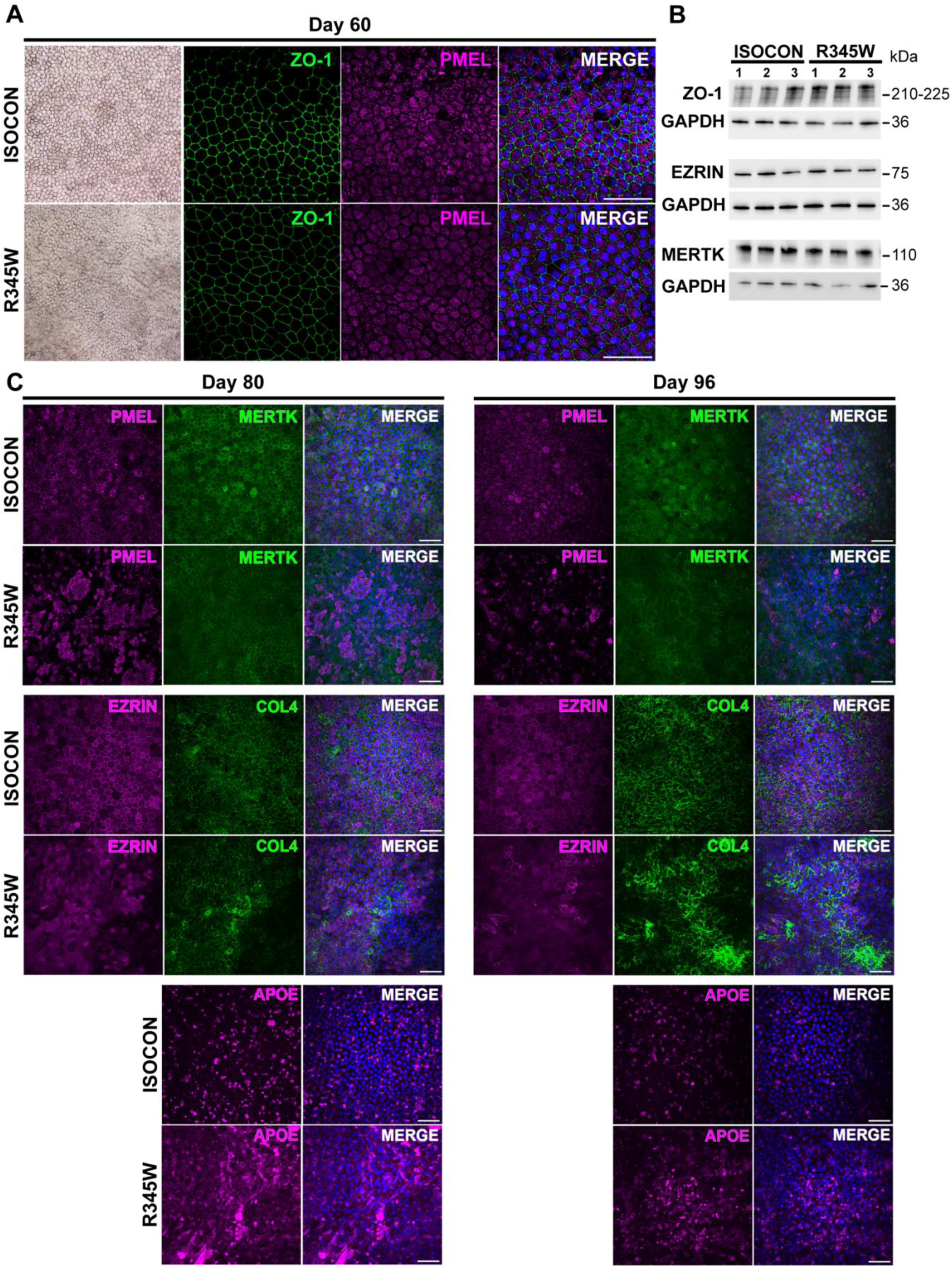
Phenotypic characterisation of isogenic and R345W iRPE. (A) Bright field microscopy and IF of iRPE from ISOCON and R345W iRPE at day 60 of differentiation, showing pigmented cobblestone morphology, tight junction formation (ZO-1, green) and pre-melanosome (PMEL, magenta) protein expression. Nuclei (blue) are labelled with DAPI. Scale bars, 50 μm. (B) Western blot analysis of protein extracts from ISOCON and R345W iRPE for ZO-1, Ezrin and MERTK. (C) IF of ISOCON and R345W iRPE at day 80 of differentiation and (D) at day 96 of differentiation. Progressive disorganisation of the R345W iRPE is evident with patchy PMEL17 (magenta) expression and Ezrin (green) disorganisation, and accumulation of drusen-associated proteins COL4 (green) and APOE (magenta). Nuclei (blue) are labelled with DAPI. Scale bars, 50 μm.

At day 80 and day 96, immunostaining of the ISOCON and R345W iRPE revealed phenotypic differences in the pattern and localisation of key RPE markers (Figure 1C). The R345W iRPE displayed a markedly patchy and heterogenous PMEL distribution, with areas of diminished signal interspersed among irregularly distributed PMEL-positive regions, potentially indicating a delay in iRPE maturity relative to the ISOCON iRPE. MERTK immunostaining was primarily localised to the plasma membrane as expected and illustrated the differences in cell morphology between the ISOCON and R345W iRPE at both timepoints, with a less organised and more heterogeneous cell population apparent in the R345W line, that deteriorated between the day 80 and day 96 timepoints. Immunostaining of collagen IV (COL4), a marker of the basement membrane, exhibited an evenly distributed and homogenous pattern, reflecting a well-organised basement membrane in ISOCON iRPE. Conversely, the COL4 immunolocalisation in the R345W iRPE, exhibited an uneven pattern, with bright accumulations, reflecting a disrupted basement membrane. Similarly, Ezrin immunolocalisation, which marks the apical microvilli, was uniform in the ISOCON line but appeared patchy and irregular in the R345W line. APOE deposits were observed in both the ISOCON and R345W iRPE at the day 80 and 96 timepoints; however, deposits were larger and more extensively distributed in the R345W line, suggesting that this substitution leads to increased APOE accumulation, a major component of drusen formation. These results collectively showed that both the ISOCON and R345W iPSC lines could form RPE, as evidenced by the expression of hallmark RPE markers. However, the presence of the R345W variant results in phenotypic alterations, suggesting that while fundamental RPE differentiation is preserved, the variant disrupts the structural organisation of the extracellular matrix components, potentially compromising RPE function and contributing to disease pathology.

### Apicobasal characteristics of patient iPSC-iRPE (iRPE)

To characterise the iRPE and confirm their apicobasal polarity, day 98 vertically embedded cross sections were immunostained for RPE markers (Figure 2). The ISOCON iRPE formed a well-organised, uniform monolayer. Ezrin was appropriately localised to the apical surface, with COL4 immunolocalisation marking a continuous and uniform basal layer (Figure 2A). In contrast, the R345W iRPE displayed a disorganised phenotype, with a dysmorphic basal membrane and the cells failing to form a uniform monolayer. COL4 immunolocalisation revealed a disrupted and thickened basement membrane, indicating abnormalities in structural integrity (Figure 2A). Distribution of PMEL localisation was comparable between the ISOCON and R345W iRPE, although ZO-1 localisation at cell-cell junctions appeared more fragmented and discontinuous in the R345W line relative to the control, potentially indicating a disruption of tight-junctions in these cells (Figure 2B). IF for EFEMP1 (fibulin 3 (F3)) was conducted in ISOCON, R345W and ISOKO iRPE (Figure 2C). ISOCON iRPE exhibited a clustered distribution of F3 across the basal layer, with the R345W iRPE also demonstrating a similar pattern but with the appearance and accumulation of more filamentous-shaped F3. Immunolocalisation of F3 in the ISOKO iRPE demonstrated both that *EFEMP1* expression was successfully knocked out of this cell line and that the antibody used was specific to F3, as no signal was observed. Moreover, COL4 immunolocalisation in the ISOKO iRPE was similar to that in the ISOCON iRPE, demarcating a continuous and well-organised basement membrane (Figure 2C).

**Figure 2.**
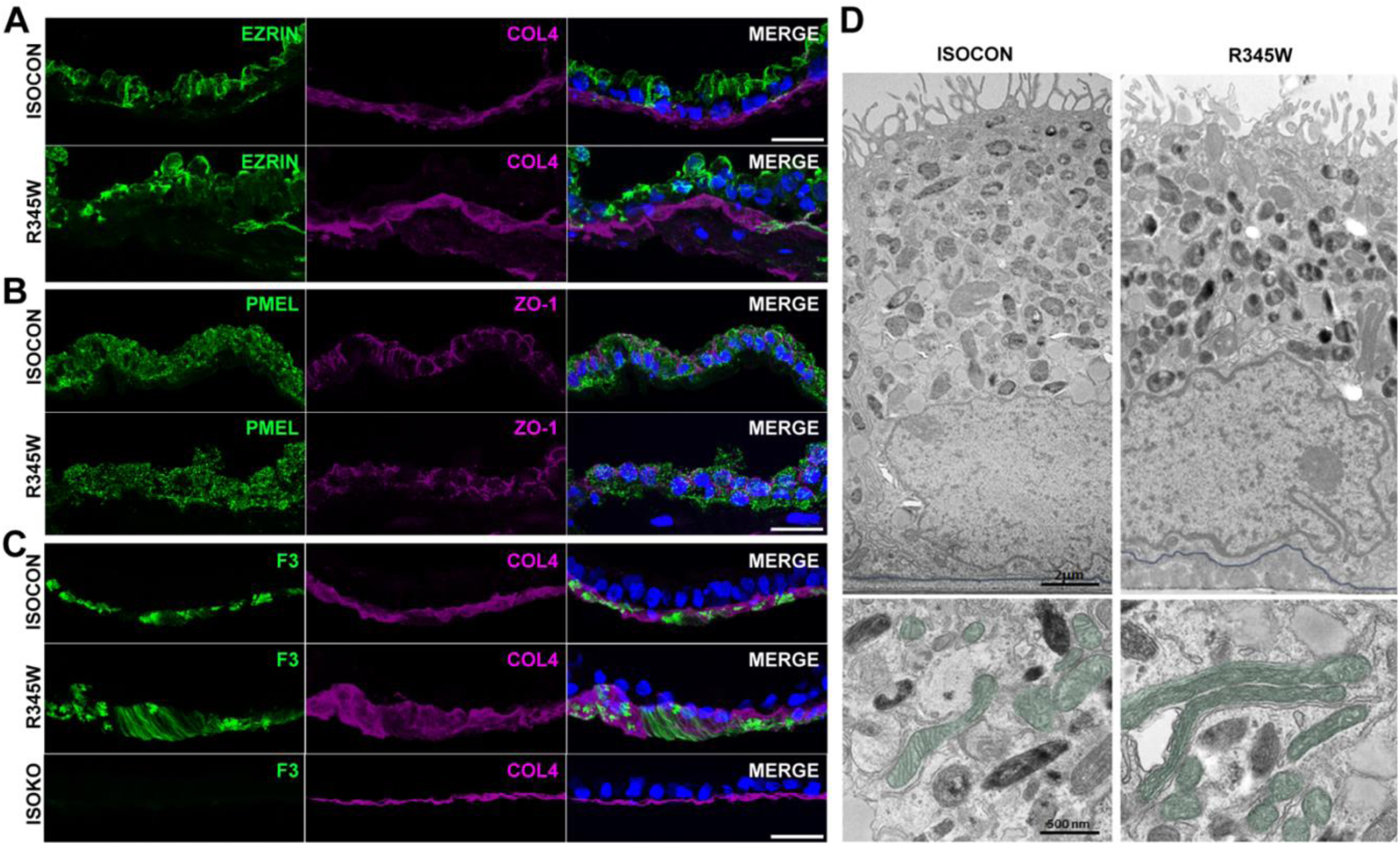
Phenotypic abnormalities are exacerbated in R345W iRPE over time. (A) IF of transverse sections (day 98 of differentiation) of ISOCON and R345W iRPE for apical marker Ezrin (green) and basement layer marker COL4 (magenta), revealing a more disorganised RPE monolayer in the R345W iRPE and remodelling of the basal layer. Nuclei (blue) are labelled with DAPI. Scale bars, 25 μm. (B) IF of transverse sections (day 98 of differentiation) of ISOCON and R345W iRPE for PMEL (green) and ZO-1 (magenta). Both ISOCON and R345W iRPE are pigmented (PMEL), but the ZO-1 staining in the R345W iRPE is irregular. Nuclei (blue) are labelled with DAPI. Scale bars, 25 μm. (C) IF of transverse sections (day 98 of differentiation) of ISOCON, R345W, and ISOKO iRPE for F3 (green) and COL4 (magenta). IF shows disorganisation of the R345W iRPE monolayer, basal accumulation of COL4 and altered organisation of F3. No accumulation of COL4 and F3 is seen in the ISOKO iRPE. Nuclei (blue) are labelled with DAPI. Scale bars, 25 μm. (D) Upper panel: Representative TEM images showing characteristic morphology of polarized pigmented RPE in ISOCON and R345W iRPE. Basal deposits are visible between the basal membrane (blue outline) and the underlying transwell membrane in R345W iRPE. Scale bar, 2 μm. Lower panel: Representative TEM images showing altered mitochondria morphology (false-coloured green) in R345W iRPE. Scale bar, 500 nm.

The ultrastructure of ISOCON and R345W iRPE was examined by TEM at day 158 of differentiation (Figure 2D). Both ISOCON and R345W iRPE demonstrated a typical polarised RPE morphology with basally located nuclei, numerous elongated microvilli extending from the apical surface and cytoplasm rich in melanin granules concentrated apically (Figure 2D, upper panel). However, basal deposits were visible between the basement membrane (blue outline) and the underlying transwell membrane in patient-derived iRPE cells (Figure 2D, upper panel). Additionally, both ISOCON and R345W iRPE contained abundant mitochondria. However, altered morphology of mitochondria (false-coloured green), which were sometimes elongated, were observed in the R345W iRPE with the cristae often appearing to be orientated longitudinally (Figure 2D, lower panel), raising the possibility of mitochondrial dysfunction as a feature of the disease pathology.

### *In vitro* screen of antisense oligonucleotides targeting *EFEMP1* c.1033C>T

Four 18-mer ASO (ASO1-4) spanning the c.1033C>T variation in *EFEMP1* (Table S4) were designed to induce RNase H1-mediated degradation of the *EFEMP1* target mRNA and analysed *in silico* as described previously.^21^ The thermodynamic binding stability of each ASO sequence was analysed *in silico* using a 124 bp sequence comprising exon 10 of R345W *EFEMP1* as target RNA (RNAstructure, University of Rochester) to determine the difference in free energy of the target sequence alone in comparison to the ASO bound target sequence, as shown for ASO1 (Figure 3A). This analysis confirmed the stable binding of all ASO to the target sequence. An *in vitro* screen of ASO1-4 in HEK293T cells was conducted to determine their ability to knock down *EFEMP1* (Figure 3B-E). HEK293T cells co-transfected with R345W *EFEMP1-*mScarlet and WT *EFEMP1*-FLAG were treated with control ASO (CTRL) (50 nM) or increasing concentrations (25 nM, 50 nM, 100 nM, 200 nM) of ASO1, 2, 3 or 4.

**Figure 3.**
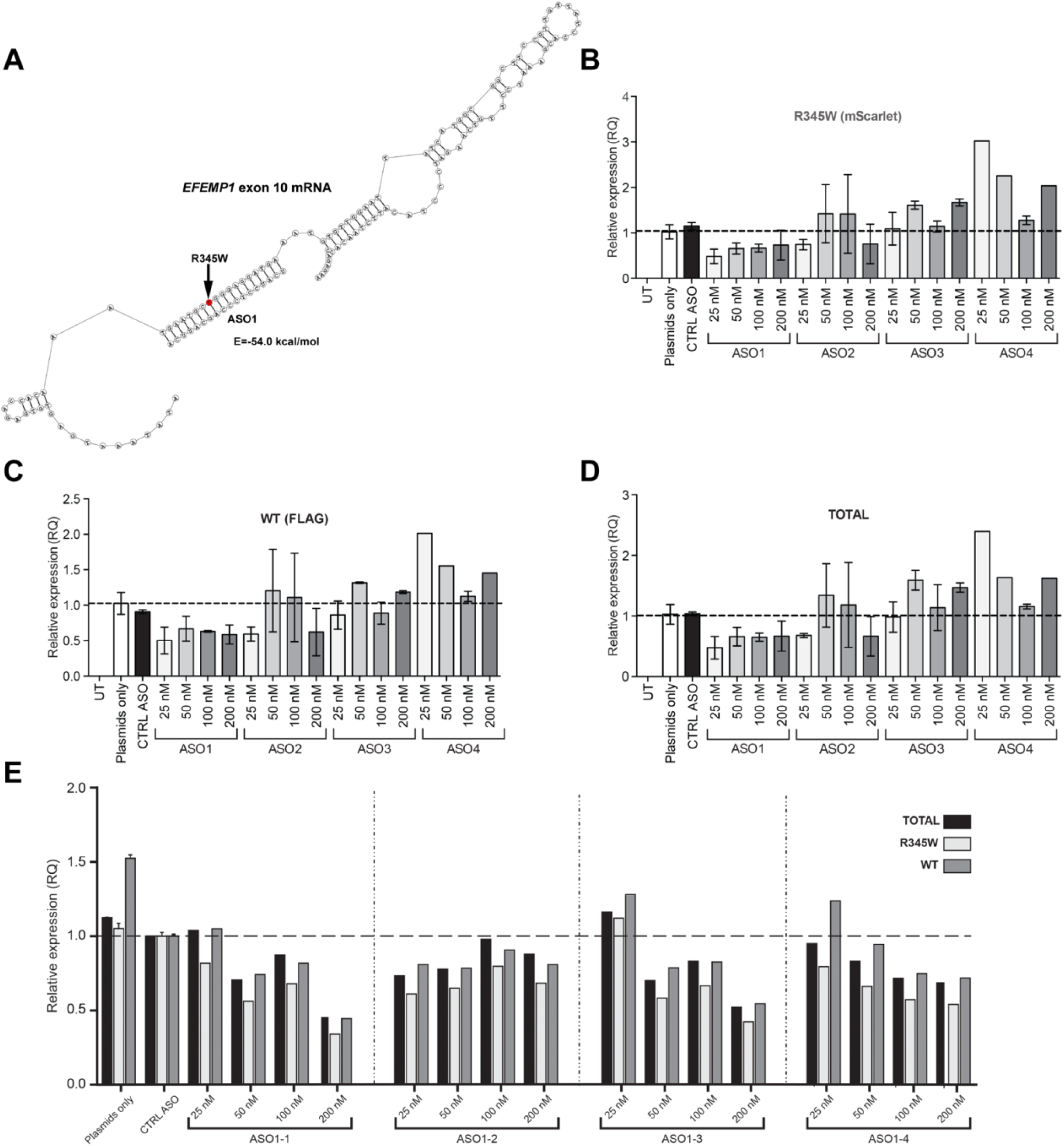
*In vitro* screen of *EFEMP1*-targeting ASO in HEK293T cells. (A) *In silico* analysis of *EFEMP1* (exon 10 mRNA) (RNAstructure University of Rochester). The c.1033C>T, p.Arg(345Trp) variant is highlighted by an arrow and the binding of ASO1 shown. The calculated difference in free energy between the target sequence alone (lowest free energy value of -21.9 kcal/mol) and the ASO1-bound target sequence (-54.0 kcal/mol) is >21 kcal/mol meeting the requirements for stable ASO targeting.^21^ (B, C, D) qPCR analysis of R345W *EFEMP1* (B), WT *EFEMP1* (C) and TOTAL *EFEMP1* (D) transcript levels following transfection of HEK293T cells with pEFEMP1(R345W)-mScarlet and pEFEMP1(WT)-3xFLAG together with control ASO (CTRL) (50 nM) or increasing amounts (25 nM, 50 nM, 100 nM, 200 nM) of ASO1, ASO2, ASO3 and ASO4. *EFEMP1* transcript levels are expressed relative to the untreated ‘plasmid only’ condition. No expression of *EFEMP1* was detected in cells that were treated with transfection reagent only (untransfected, UT). The experiment was repeated three times (N=3) with four technical replicates (n=4) per condition in each experiment. Bars represent mean ± SEM. (E) qPCR analysis of TOTAL (black bars), R345W (light grey bars) and WT (dark grey bars) *EFEMP1* transcript levels following the transfection of HEK293T cells with pEFEMP1(R345W)-mScarlet and pEFEMP1(WT)-3xFLAG together with control ASO (CTRL) (50 nM) or increasing amounts (25 nM, 50 nM, 100 nM, 200 nM) of ASO1.1, ASO1.2, ASO1.3 and ASO1.4. *EFEMP1* transcript levels are expressed relative to the untreated ‘plasmid only’ condition. The experiment was repeated three times (N=3) with four technical replicates (n=4) per condition in each experiment. Bars represent mean ± SD.

Quantitation of R345W *EFEMP1* (Figure 3B), WT *EFEMP1* (Figure 3C) and total *EFEMP1* (Figure 3D) transcripts by qPCR showed a reduction in all *EFEMP1* levels (R345W, WT, TOTAL) compared to the CTRL ASO and untreated cells (plasmids only) only with ASO1 (Figure 3). Reduced levels of *EFEMP1* (R345W, WT, TOTAL) with ASO2 were evident only at the highest and lowest concentrations. No reduction of *EFEMP1* R345W, WT or TOTAL transcript levels was evident with ASO3 or with ASO4. No expression of *EFEMP1* was detected in cells that were mock transfected (UT). Four derivatives of the best-preforming ASO1 (ASO1.1-1.4) (Table S4) were tested *in vitro* in HEK293T cells (Figure 3E). Levels of *EFEMP1* (TOTAL, R345W, WT) were measured by qPCR. All ASO, except for ASO1.2, showed a concentration-dependent decline in *EFEMP1* (TOTAL, R345W, WT) transcript levels compared to the untreated (plasmid only) and control ASO (CTRL) conditions, and the highest level of *EFEMP1* reduction was achieved with 200 nM ASO1.1. Whilst ASO1.2 could effectively reduce *EFEMP1* levels at all concentrations tested, the ASO1.2-mediated reduction in *EFEMP1* levels was not concentration-dependent. All ASO (1.1-1.4) mediated a greater reduction in R345W *EFEMP1* compared to WT *EFEMP1* (Figure 3E).

### Allele-specific *EFEMP1* targeting following ASO transient transfection of iRPE

To assess the efficacy of ASO treatment in the downregulation of *EFEMP1* transcript expression, ASOs 1.1-1.4 were transiently transfected into R345W EFEMP1 patient-derived iRPE at a dose of 200 nm at day 72 of differentiation. Transfection of the 6-FAM-labelled ASO1.1 into the R345W iRPE, resulted in strong intracellular fluorescence (Figure 4A). Experimental cell samples were lysed 48 hours post-transfection for transcript-level analyses. qPCR analysis demonstrated a significant 30-40% reduction of total *EFEMP1* transcript levels in therapeutic ASO-treated conditions, relative to transfection reagent only (NT) and control ASO treated conditions (CTRL), suggesting an effective downregulation of *EFEMP1* expression by the designed ASOs (Figure 4B).

**Figure 4.**
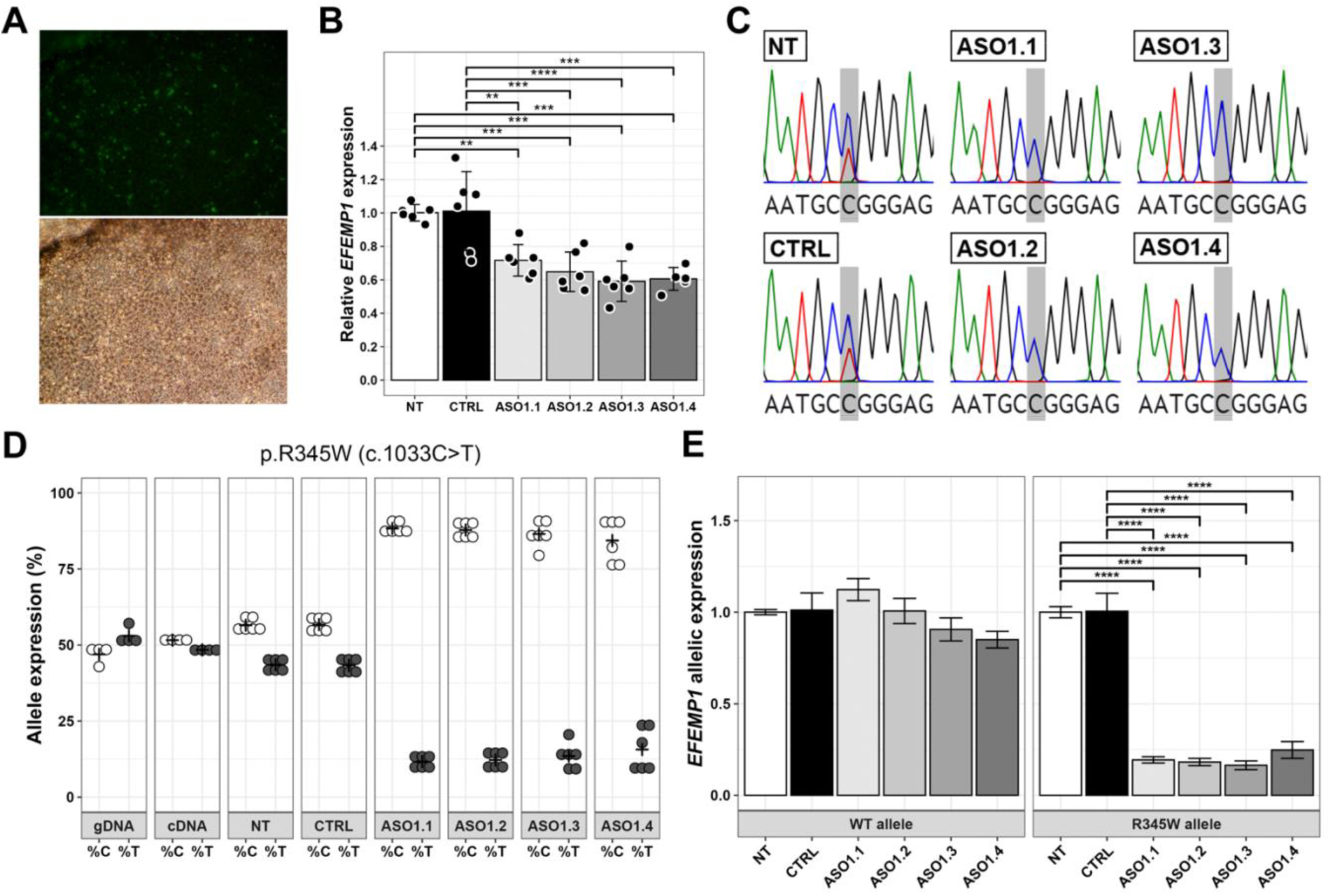
*EFEMP1* allele-specific targeting in iRPE following ASO transfection. (A-D) iRPE were transfected with ASO1.1, 1.2, 1.3, 1.4 or control ASO (CTRL) (200 nM) at day 72 of differentiation and analysed 48 hours later. (A) Fluorescence and bright field images of R345W iRPE showing efficient delivery of 6-FAM conjugated ASO. (B) Quantitation of *EFEMP1* transcript levels by qPCR. All ASO mediate a significant decrease in *EFEMP1* levels. N = 3 independent experiments, n = 2 technical replicates of each condition per experiment. Statistical significance was determined by one-way ANOVA followed by post-hoc Tukey’s (HSD) test where *, ** and *** denotes a *p* value <0.05, 0.01 and 0.005 respectively. ns = not significant. Bars represent mean ± SEM. (C) Sanger sequencing chromatograms of untreated (NT) R345W iRPE and R345W iRPE treated with CTRL ASO, ASO1.1, ASO1.2, ASO1.3 or ASO1.4 (200 nM). The heterozygous ‘C/T’ peak in the NT and CTRL treated R345W iRPE is resolved to a homozygous ‘C’ peak in all ASO treated samples. (D) Quantitation of ASO-mediated allele-specific *EFEMP1* targeting by NGS. WT (%C) and R345W (%T) *EFEMP1* transcript levels were quantified in untreated R345W iRPE (NT) or R345W iRPE treated with CTRL or therapeutic ASO1.1, 1.2, 1.3 or 1.4. NGS results for gDNA and cDNA from corresponding patient renal epithelial cells are also shown. N = 3 independent experiments, n = 2 technical replicates of each condition per experiment. Bars represent mean ± SD. (E) Integrated analysis of the qPCR and NGS data showing a significant reduction of the R345W transcript with all ASO. Statistical significance was determined by one-way ANOVA followed by post-hoc Tukey’s (HSD) test where *, ** and *** denotes a *p* value <0.05, 0.01 and 0.005 respectively. ns = not significant. Bars represent mean ± SD.

To explore allele-specificity, Sanger sequencing of cDNA amplicons spanning the c.1033C>T variant revealed preferential downregulation of the R345W transcript (Figure 4C). Electropherogram traces from therapeutic ASO-treated samples demonstrated marked reduction in the peak corresponding to the variant ‘T’ base relative to NT and CTRL conditions, providing semi-quantitative confirmation of allele-specificity of the ASOs.

Targeted next generation sequencing (NGS) was conducted using primers (Table S5) producing an amplicon covering the c.1033C>T locus of *EFEMP1* to enable quantitation of the allelic discrimination of the ASOs (Figure 4D). A baseline of allelic expression in the DHRD patient-derived samples was obtained to observe the possibility of *EFEMP1* allele skewing (Figure 4D). Amplicon quantitation of gDNA from patient-derived renal epithelial cells confirmed the expected 50:50 ratio between the WT and R345W alleles of *EFEMP1*, confirming equal copy number. cDNA prepared from renal epithelial cells of the patient also demonstrated a 50:50 ratio between the WT and R345W alleles of *EFEMP1*, indicating that in this cell type, both alleles of *EFEMP1* are equally expressed. However, basal expression levels of the cDNA prepared from differentiated iRPE in the NT and CTRL ASO treated conditions revealed a 55:45 WT:R345W allelic expression split, indicating that the WT allele is expressed slightly higher than the R345W allele in differentiated iRPE. Following therapeutic ASO treatment, a clear reduction in R345W allele expression was observed for all ASOs, corroborating the Sanger sequencing data (Figure 4C). Therapeutic ASO treatment through transient transfection resulted in the WT allele constituting 84.4-88.3% of total allelic expression, and the R345W allele constituting 11.7-15.6% of total allelic expression (Figure 4D).

Integrated analysis of the qPCR (Figure 4B) and NGS (Figure 4D) was performed to gain an understanding of how the ASO treatments affect the expression of the *EFEMP1* WT and R345W alleles relative to total *EFEMP1* transcript expression (Figure 4E). The percentage of allelic expression was calculated in comparison to total levels of *EFEMP1* transcript (Figure 4B) before normalisation of the data to the NT control condition for the respective data group (WT or R345W allele) (Figure 4D). No significant differences between the control (NT and CTRL ASO) and therapeutic ASO treatment conditions were observed for the expression of the WT allele, with significant reductions in the R345W allele expression clearly observed following therapeutic ASO treatment. These results thus demonstrated consistent R345W allele-specific downregulation following treatment with the therapeutic ASOs. Based on these data, ASO1.1 was taken forward for the rest of the ASO treatments, due to its high reproducibility in reducing R345W *EFEMP1* allele expression and its ideal length (16-mer).

### Allele-specific *EFEMP1* targeting following ASO gymnosis of iRPE

Gymnotic (unassisted) delivery of ASO1.1 at day 60 of differentiation was carried out to evaluate ASO efficacy in a setting that more closely recapitulates physiological uptake mechanisms (Figure 5). Fluorescence microscopy of gymnotically delivered 6-FAM-conjugated ASO1.1 (1 μM) showed efficient uptake across the R345W iRPE population following a 7-day treatment (Figure 5A). The ASO was demonstrated to robustly accumulate in the nucleus, with additional punctate localisation in the cytoplasm. This distribution is consistent with the known subcellular activity of RNase H1, which is present in both the nucleus and cytoplasm.^22^ While the ASO is designed to target mature mRNA, their nuclear localisation suggests potential additional engagement with pre-mRNA targets. RNase H-mediated cleavage of the target transcript can occur in both subcellular compartments, with cytoplasmic ASO signal potentially indicating interactions with mature transcripts or transient endosomal retention. Thus, ASO localisation in both of these compartments indicates enhanced likelihood of effective target knockdown following gymnotic uptake.

**Figure 5.**
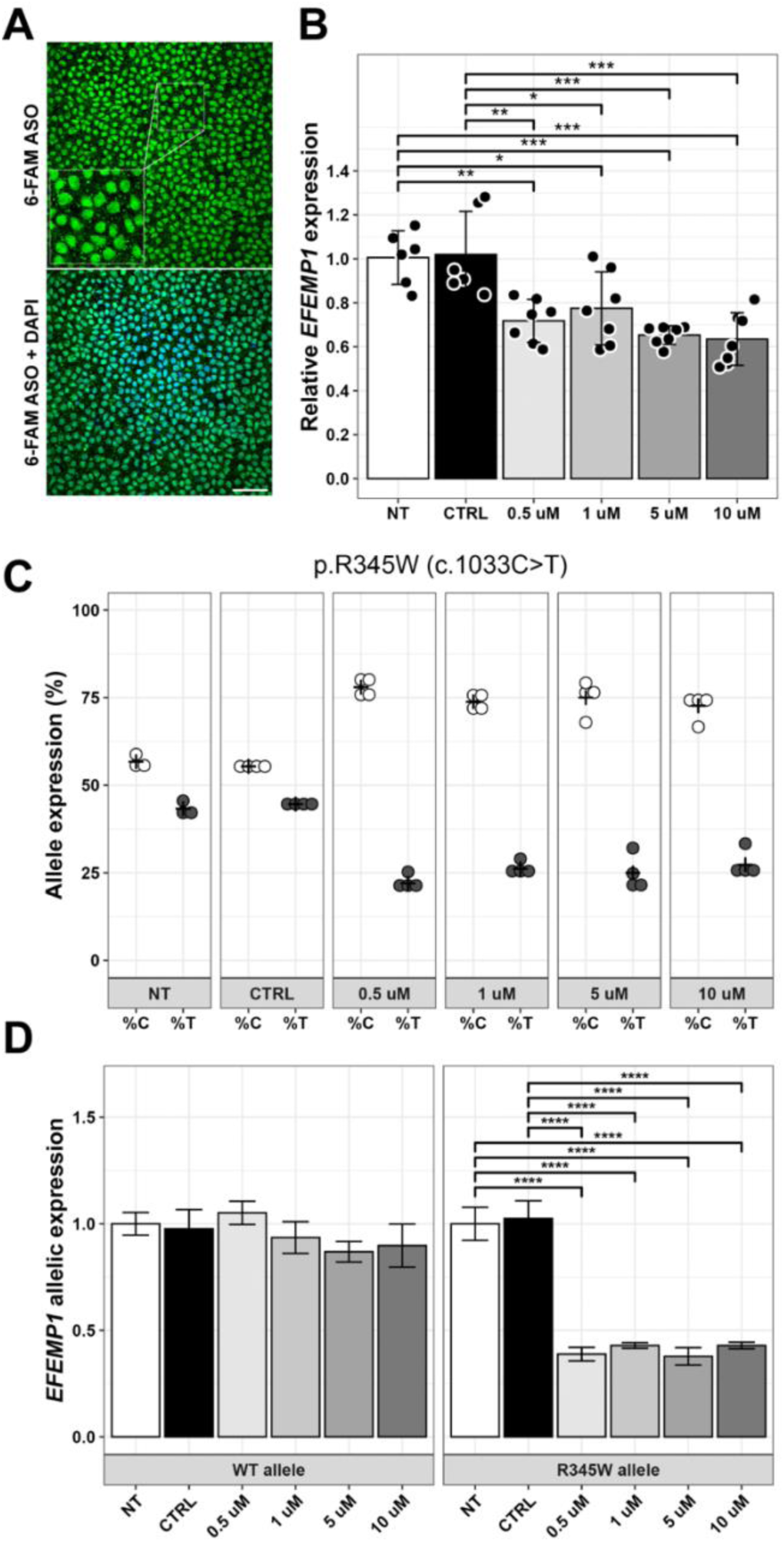
*EFEMP1* allele-specific targeting in iRPE following ASO gymnosis. (A-C) R345W iRPE at day 60 of differentiation underwent a 7-day treatment with control ASO (CTRL) (1 μM) or ASO1.1 at the doses 0.5 µM, 1 µM, 5 µM and 10 μM. (A) Fluorescence from gymnotically delivered 6-FAM-conjugated ASO1.1 (1 μM) showed efficient uptake predominately in the nucleus and extra-nuclear endocytic vesicles. Nuclei are labelled with DAPI. Inset shows higher magnification image. Scale bar, 50 μm. (B) Quantitation of *EFEMP1* transcript levels by qPCR. All doses of ASO1.1 mediate a significant decrease in *EFEMP1* expression levels relative to the non-treated (NT) and CTRL ASO-treated R345W iRPE. N = 2 independent experiments, n = 3 to 4 technical replicates for each condition per experiment. Statistical significance was determined by one-way ANOVA followed by post-hoc Tukey’s (HSD) test where *, ** and *** denotes a p value <0.05, 0.01 and 0.005 respectively. Bars = mean ± SD. (C) Quantitation of ASO1.1-mediated allele-specific *EFEMP1* targeting by NGS at doses 0.5 µM, 1 µM, 5 µM and 10 μM. WT (%C) and R345W (%T) *EFEMP1* transcript levels were quantified in untreated R345W iRPE (NT) or R345W iRPE treated with CTRL or ASO1.1. N = 2 independent experiments, n = 2 technical replicates for each condition per experiment. (E) Integrated analysis of the qPCR and NGS data showing a significant reduction of the R345W transcript with all ASO1.1 doses. Statistical significance was determined by one-way ANOVA followed by post-hoc Tukey’s (HSD) test where *, ** and *** denotes a p value <0.05, 0.01 and 0.005 respectively. Bars = mean ± SD.

R345W iRPE underwent a 7-day treatment with ASO1.1 at the doses 0.5 µM, 1 µM, 5 µM and 10 µM, and *EFEMP1* gene expression was analysed by qPCR (Figure 5B). Therapeutic ASO treatments at each of the doses led to a significant downregulation of total *EFEMP1* expression relative to the NT and CTRL ASO treatment conditions. However, no clear dose-dependent response was observed between therapeutic ASO treatments. The magnitude of *EFEMP1* knockdown remained comparable between the lowest and highest ASO doses tested. This plateau in response indicates that maximal target suppression could be achieved at or below the lowest tested concentration.

Next, targeted NGS was employed to evaluate the allelic discrimination of gymnotically delivered ASOs (Figure 5C). Baseline allelic expression in iRPE for the DHRD patient included in this study was confirmed at a 55:45 WT:R345W allelic split, as observed in the NT and CTRL ASO conditions. Again, there was no dose-dependent response following ASO1.1 treatment at the doses 0.5 µM, 1 µM, 5 µM and 10 µM as all doses led to the WT allele constituting 73-78% and the mutant allele constituting 22-27% of total *EFEMP1* expression.

Integrated analysis of the qPCR (Figure 5B) and targeted NGS (Figure 5C) data was carried out to explore the efficacy of ASO1.1 in its allele-specific downregulation of *EFEMP1*. The percentage of allelic expression was calculated in comparison to total levels of *EFEMP1* transcript (Figure 5B) with normalisation of the data to the NT control condition for the respective data group (WT or R345W allele) (Figure 5D). Similar to the qPCR and NGS data collation carried out for the transient transfection (Figure 4D), no significant differences between the controls (NT and CTRL ASO) and therapeutic ASO treatment conditions were observed for WT allele expression, whereas significant reductions in mutant *EFEMP1* was clearly observed. Overall, this confirmed consistent *EFEMP1* R345W allele-specific downregulation via ASO1.1 gymnosis.

### Rescue of patient iRPE phenotype following ASO gymnosis

We next examined the phenotype of patient iRPE following gymnotic treatment with the therapeutic ASO1.1 (Figure 6). To assess the impact of ASO gymnosis on APOE, whole-mount iRPE samples following a 1-week treatment starting from day 60 of differentiation were immunolabelled for APOE (Figure 6A). APOE has been shown to accumulate in sub-RPE deposits and contribute to drusen formation, hallmarks of DHRD. IF analysis revealed low APOE signal for both the ISOCON and ISOKO iRPE. In contrast, NT R345W iRPE exhibited a marked increase in APOE fluorescence, consistent with the DHRD disease phenotype. 1-week gymnosis of ASO1.1 at a dose of 1 µM resulted in a substantial reduction in APOE fluorescence, as illustrated in the representative images and by quantification of total fluorescence intensity (Figure 6A and B). Quantitative analysis confirmed a significant elevation of APOE levels in the R345W NT condition relative to ISOCON and ISOKO iRPE. There was no difference in the levels of APOE quantified in the ISOCON and ISOKO iRPE. APOE levels in the ASO-treated R345W iRPE displayed significantly reduced APOE fluorescence relative to the NT condition, although expression remained modestly higher than in the ISOCON and ISOKO condition, following the 1-week treatment. These findings indicate that suppression of R345W *EFEMP1* via gymnotically delivered ASO mitigates downstream APOE accumulation, implicating a functional relationship between *EFEMP1*-driven pathology and APOE accumulation in this iRPE model.

**Figure 6.**
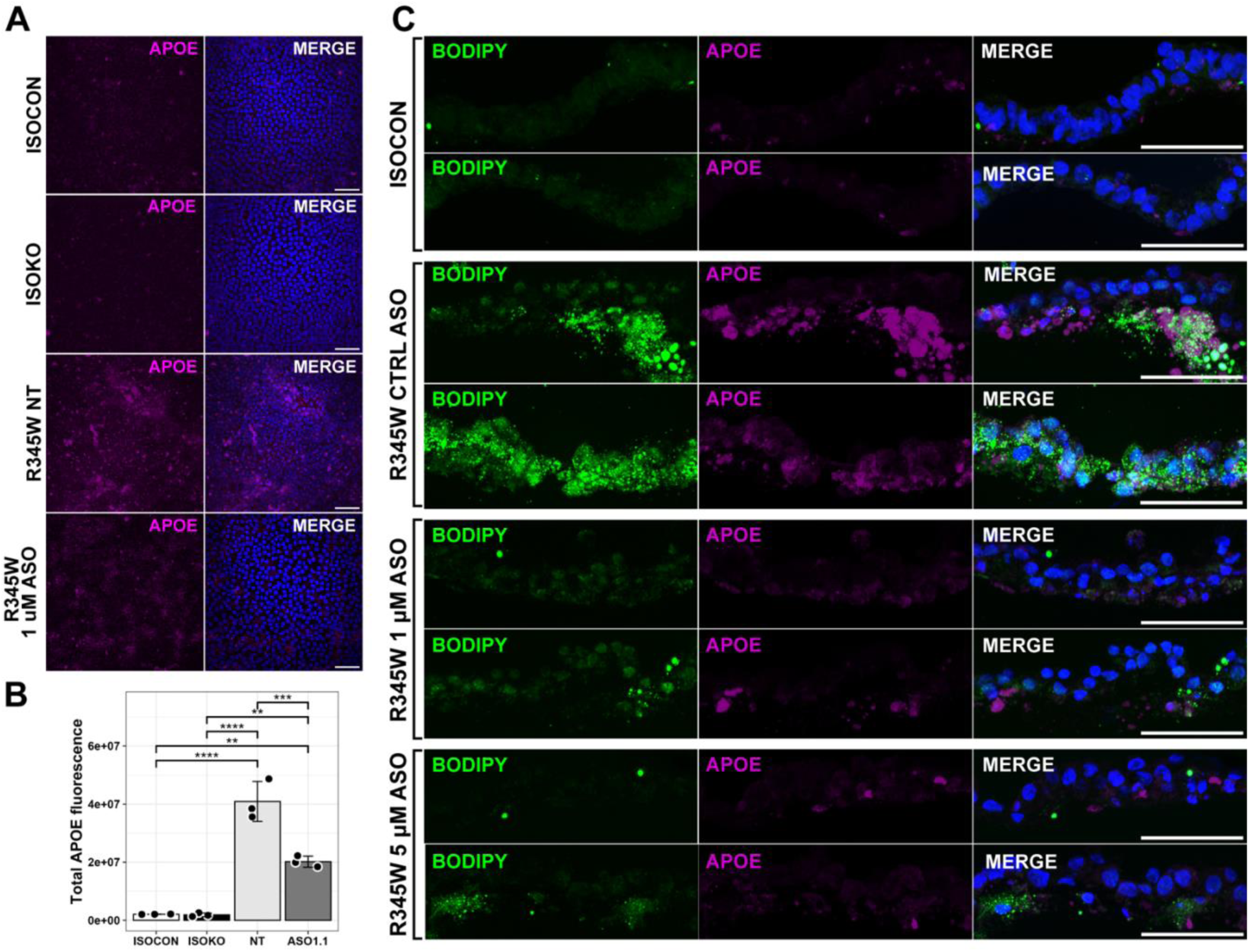
Phenotypic rescue of APOE and lipid accumulation in R345W iRPE following ASO gymnosis. (A) IF of APOE in ISOCON, ISOKO, untreated R345W (NT) and R345W iRPE treated gymnotically with ASO1.1 (1 μM dose for 7 days from day 60 of differentiation), showing APOE (magenta) accumulation in R345W iRPE and rescue in ASO1.1 treated R345W iRPE. Nuclei are labelled with DAPI. Scale bars, 50 μm. (B) Quantitation of APOE (relative fluorescence levels) in ISOCON, untreated R345W iRPE (NT) and ASO1.1 treated R345W iRPE (1 μM dose for 7 days from day 60 of differentiation). Fluorescence levels were quantified from four independent images. Statistical significance was determined by one-way ANOVA followed by post-hoc Tukey’s (HSD) test where *, ** and *** denotes a p value <0.05, 0.01 and 0.005 respectively. Bars = mean ± SD. (C) Images of transverse sections (14 μM) from ISOCON and R345W iRPE treated gymnotically with CTRL ASO or with therapeutic ASO1.1 (1 μM and 5 μM dose for 14 days from day 84 of differentiation), showing accumulation of neutral lipid (BODIPY, green) and basal APOE (magenta) accumulation in R345W iRPE that is rescued following ASO1.1 treatment. Nuclei are labelled with DAPI. Scale bars, 20 μm.

Next, the distribution of APOE and lipid accumulation in vertically embedded iRPE cross sections was assessed following a 2-week gymnotic ASO1.1 treatment, initiated on day 84 of differentiation, at concentrations of 1 µM and 5 µM (Figure 6C). Sections were stained with BODIPY to visualise neutral lipid content and co-labelled for APOE. ISOCON iRPE exhibited minimal BODIPY and APOE immunoreactivity, whereas patient-derived iRPE treated with CTRL ASO displayed elevated fluorescence signals for both markers. In contrast, ASO1.1-treated iRPE showed a pronounced reduction in BODIPY and APOE fluorescence intensity at both 1 μM and 5 μM doses, indicating that gymnotic ASO1.1 administration effectively decreases APOE expression and lipid accumulation in patient-derived DHRD iRPE within two weeks.

To further investigate the robustness of ASO1.1 effects and its potential impact on ECM organisation, a 2-month gymnotic treatment was conducted on patient-derived iRPE (Figure 7). ISOCON, NT R345W and 1 µM ASO-treated R345W iRPE cultured on chamber slides were analysed for BODIPY, APOE, COL4 and F3 expression and localisation. Consistent with our earlier findings, ISCOCON iRPE show minimal BODIPY and APOE signal (Figure 7A), indicating low lipid and APOE accumulation. Conversely, the R345W NT condition exhibited strong immunostaining for both markers. ASO1.1 treatment (1 μM) in R345W iRPE significantly reduced both BODIPY and APOE fluorescence (Figure 7A, C, D). IF was also conducted with antibodies to COL4 and F3 (Figure 7B). ISOCON iRPE showed a homogenous distribution of COL4 staining, in contrast to uneven patches of accumulated COL4 observed in the NT R345W condition (Figure 7B). ASO1.1 treatment significantly attenuated COL4 accumulation, resulting in a more even staining pattern (Figure 7B, E), supporting a potential role for mitigating ECM disorganisation induced by R345W *EFEMP1* expression. These findings indicate that reduction of R345W *EFEMP1* expression not only attenuates lipid and APOE accumulation, but also promotes the rescue of pathological ECM features in DHRD iRPE.

**Figure 7.**
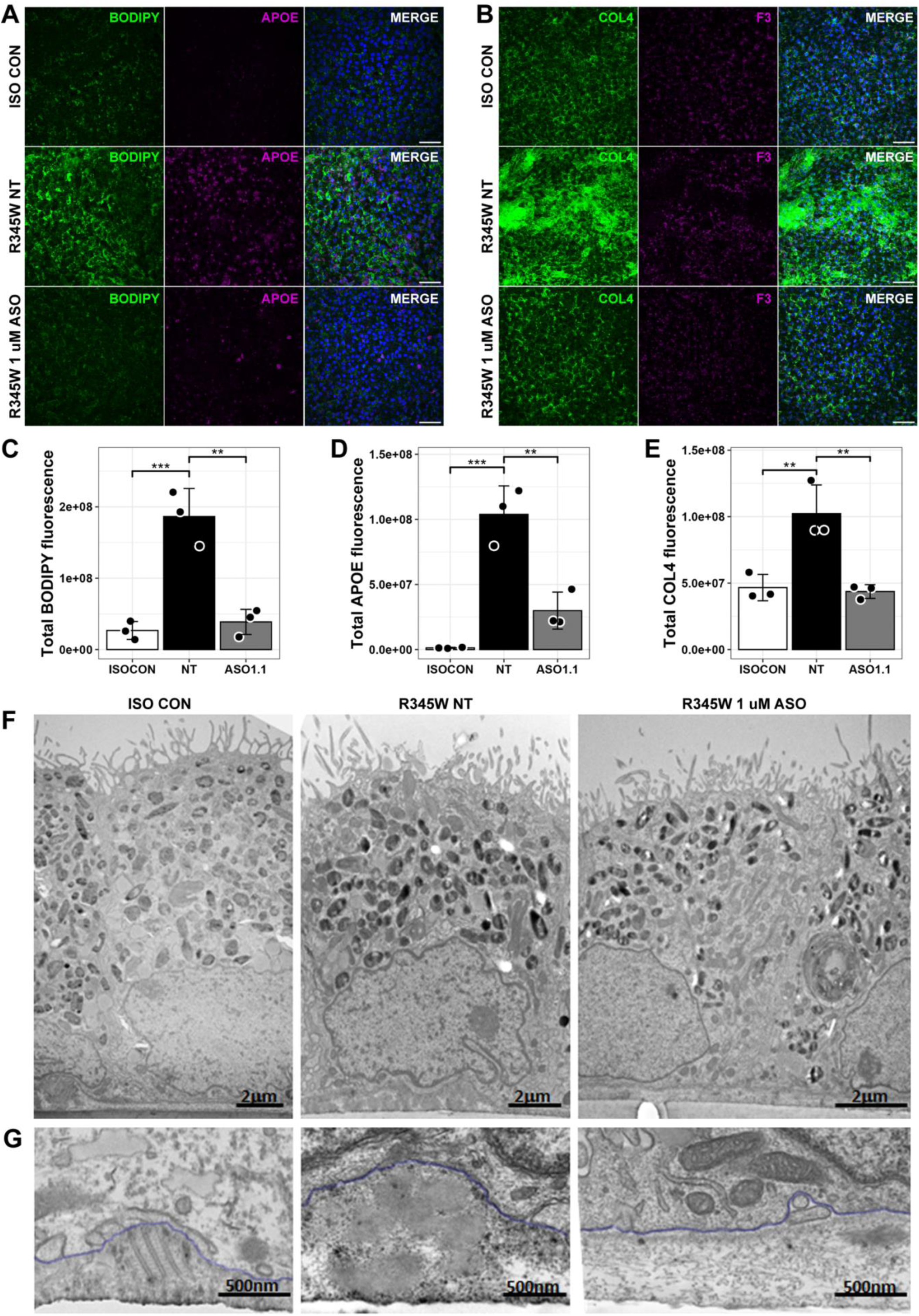
Phenotypic rescue of APOE, ECM abnormalities and sub-RPE lipid deposition in R345W iRPE following ASO gymnosis. (A, B) Staining of ISOCON, untreated R345W (NT) and R345W iRPE following 2-month ASO1.1 gymnosis (1 μM) from day 100 of differentiation. (A) Accumulation of lipids (BODIPY, green) and APOE (magenta) was evident in NT R345W iRPE and was rescued following treatment with therapeutic ASO1.1. Nuclei are labelled with DAPI. Scale bars, 50 μm. (B) Accumulation of COL4 and F3 is evident in NT R345W iRPE and is rescued following treatment with therapeutic ASO1.1. Nuclei are labelled with DAPI. Scale bars, 50 μm. (C-E) Quantitation of relative fluorescence levels of BODIPY (C), APOE (D) and COL4 (E). Statistical significance was determined by one-way ANOVA followed by post-hoc Tukey’s (HSD) test where *, ** and *** denotes a p value <0.05, 0.01 and 0.005 respectively. Bars = mean ± SD. (F) TEM of ISOCON, untreated R345W (NT) and R345W iRPE following 2-month ASO1.1 gymnosis (1 μM) from day 100 of differentiation. Upper panel: ISOCON, untreated R345W and ASO1.1 treated R345W iRPE show a characteristic RPE-like morphology. Scale bars, 2μm. Lower panel: Representative TEM images of basal deposits between the basal membrane (blue outline) and the underlying transwell membrane. Scale bars, 500 nm.

Finally, TEM of ISOCON, NT R345W and ASO1.1-treated R345W iRPE was conducted at day 158 following 2-month ASO1.1 gymnosis (1 μM) commencing at day 100 of differentiation. All cells showed characteristic RPE-like morphology, forming monolayers of polarized pigmented cells (Figure 7F). Accumulation of microfibrils (blue outline) was observed in the ISOCON iRPE between the basal membrane and underlying transwell membrane (24% of surface) (Figure 7G). In contrast, lipid-rich deposits (blue outline) were observed in the NT R345W-iPSC (22% of surface) with microfibril accumulation in the extracellular matrix observed only rarely (5% of surface) (Figure 7G). Lipid-rich deposits were almost completely absent (3% of surface) from R345W iRPE following 2-month gymnosis with ASO1.1 (1 μM) (Figure 7G).

## DISCUSSION

In this study, we confirm that both patient-derived and isogenic iRPE initially exhibit a typical RPE-like morphology and express key RPE markers associated with epithelial identity and polarity, tight junction formation and melanosome biogenesis. However, the c.1033C>T, p.Arg(345Trp) variant results in progressive phenotypic alterations in the patient-derived iRPE. While the corrected isogenic iRPE formed a well-organised, uniform monolayer, the patient-derived iRPE fail to form a uniform monolayer and become progressively less well-organised, with tight junctions appearing more fragmented and discontinuous. Moreover, the basement membrane in the isogenic control iRPE is well-organised, forming a continuous and uniform basal layer. In contrast, a dysmorphic basal layer characterised by progressive disruption and structural disorganisation of the extracellular matrix and thickening of the basement membrane is evident in the patient-derived iRPE, indicating abnormalities in the structural integrity of this layer. Interestingly, EFEMP1 itself is localised across the basement membrane in both isogenic and patient-derived iRPE. However, in patient-derived iRPE, EFEMP1 accumulated in the basement membrane and presented a more filamentous appearance. This is consistent with the accumulation of EFEMP1 seen in the basal layer between RPE and drusen in human DHRD donor eyes, the formation of basal deposits in primary mouse RPE cells carrying the R345W *Efemp1* substitution and with the formation of sub-RPE deposits in R345W *Efemp1* knock-in mice.^7,8,11^ Finally, the accumulation of lipids and APOE, a key component of drusen, is evident in the patient-derived iRPE. While lipid-rich deposits underneath the RPE in the basal layer were seldom observed in the isogenic iRPE, increased sub-RPE lipid-rich deposits were observed in the patient-derived iRPE. Similar phenotypic characteristics have been reported in other models of DHRD using patient-derived iRPE and isogenic controls.^13,14^ Galloway *et al*. (2017) reported an increased accumulation of lipids and APOE-positive basal deposits beneath the RPE in patient-derived iRPE, in addition to increased deposition and levels of extracellular matrix and basement membrane constituents.^13^ Similarly, Tsai *et al*. (2021) reported the intracellular and extracellular accumulation of lipids in patient-derived iRPE, in addition to the downregulation of carboxylesterase 1 (CES1) and the attenuation of cholesterol efflux from these cells.^14^ The authors proposed that a hyper-inhibitory effect of R345W EFEMP1 on epidermal growth factor receptor (EGFR) signaling may suppress CES1 expression, thereby attenuating cholesterol export and leading to lipid accumulation.

In addition to the phenotypic alterations described above, altered mitochondrial morphology is a novel phenotype observed in the patient-derived iRPE, including a population of elongated mitochondria and for some mitochondria, a longitudinal orientation of the cristae. Mitochondrial dysfunction, together with oxidative stress and inflammation, is increased in age-related macular degeneration (AMD),^23–25^ but has not been previously reported in juvenile macular dystrophy, raising the possibility of a direct role for mitochondrial dysfunction in the disease pathology of these disorders. Another novel finding in our study was the occurrence of microfibrils in the extracellular matrix beneath the basement membrane in the isogenic iRPE which was largely absent from the patient-derived iRPE. The ultrastructural appearance and banded pattern of these microfibrils in the extracellular matrix suggest that they may be comprised of collagen VI.^26^ While the identity of these microfibrils in the extracellular matrix requires further scrutiny, it has been reported that ARPE-19 cells engineered to be homozygous for the R345W *EFEMP1* variant developed changes in the extracellular matrix that included a reduced and disorganised network of stretched collagen VI fibers in comparison to wild type ARPE-19, which exhibited a continuous network of fibers with some thickened and localized areas.^27^ This was accompanied by a thickened and multilayered fibronectin network in the R345W ARPE-19 cells.^27^ Overall, the data are consistent with structural disorganisation of the extracellular matrix in cellular models of DHRD.

Interestingly, the isogenic *EFEMP1* knockout iRPE in our study did not emulate the phenotypic characteristics observed in the patient-derived iRPE. The *EFEMP1* knockout iRPE formed a well-organised monolayer with a continuous and uniform basement membrane indicating maintenance of the structural integrity of the extracellular matrix. Moreover, APOE accumulation was not observed in the *EFEMP1* knockout iRPE. Although this finding cannot fully rule out subtle or undetected changes in the RPE basal layer and extracellular matrix, it does confirm the dominant toxic gain-of-function of R345W EFEMP1 in the RPE in addition to the absence of *EFEMP1* loss-of-function or haploinsufficiency as a disease mechanism underpinning DHRD pathology.

Given the toxic gain-of-function of R345W EFEMP1, we designed an antisense oligonucleotide to specifically promote the downregulation of the R345W *EFEMP1* transcript through RNase H dependent degradation. Both assisted delivery of the lead therapeutic ASO at a dose of 200 nM and gymnotic delivery over a dose range of 0.5 μM to 10 μM resulted in allele-specific targeting of the R345W *EFEMP1* transcript in the patient-derived iRPE. ASO treatment of patient-derived iRPE through transient transfection resulted in the reduction of R345W *EFEMP1* transcript levels to 11.7-15.6% of total allelic expression as quantified by next generation sequencing, whilst gymnotic treatment was less effective, reducing R345W *EFEMP1* transcript levels to 22-27% of total levels. A dose-dependent response was not observed following gymnotic delivery of the lead ASO, with the magnitude of knockdown of the R345W *EFEMP1* transcript comparable between the lowest and highest ASO doses tested, indicating that maximal target suppression could be achieved at or below the lowest tested concentration. Despite the reduced efficacy of the lead ASO to reduce the R345W *EFEMP1* transcript following gymnotic delivery, treatment of patient-derived iRPE with the lead ASO was able to effectively rescue the disease phenotype even when treatment was initiated after the onset of the phenotypic changes and within a short treatment timeframe. A one-week gymnotic treatment with ASO resulted in substantial reduction in drusen-associated APOE accumulation. Similarly, a two-week gymnotic ASO treatment administered after progression of the phenotypic changes in the patient-derived iRPE resulted in a pronounced reduction not only of APOE, but also lipid accumulation. Finally, a two-month gymnotic treatment in aged patient-derived iRPE was shown to not only attenuate APOE and sub-RPE lipid deposits but also rescue the disorganisation of the extracellular matrix. These results suggest that administration of ASO treatment after the onset of drusen formation in DHRD patients could be therapeutically beneficial in ameliorating the pathological features of the disease.

While no therapeutic oligonucleotide has been shown to be effective for the treatment of DHRD to date, AMD has been demonstrated to be amenable to administration of oligonucleotide therapies via intravitreal administration. Pegaptanib (MacugenTM) and avacincaptad pegol (IzervayTM) are small oligonucleotide RNA aptamers approved for the treatment of AMD.^28^ Macugen is an anti-angiogenic therapy targeting vascular endothelial growth factor (VEGF) for the treatment of neovascular AMD granted FDA approval in 2004 and EMA approval in 2006, while Izervay targets complement C5 for the treatment of geographic atrophy secondary to AMD and was approved by the FDA in 2023. The biological mechanisms of these aptamers differ from that of the antisense oligonucleotide investigated in this study. Macugen blocks the biological activity of VEGF by binding selectively to the exon 7-encoded heparin-binding domain of the VEGF-165 isoform thereby preventing interaction with its receptors. Izervay targets the complement system by blocking the cleavage of the C5 complement protein into C5a and C5b preventing the assembly of the membrane attack complex (MAC). The route of aptamer administration for these approved therapies includes intravitreal injection or scleral injection via the inferotemporal quadrant. Several other aptamers are currently under clinical investigation for the treatment of AMD, including E10030 (Fovista®), umedaptanib pegol (APT-F2P, RBM-007) and AS1411.^28^ Moreover, several small interfering RNA (siRNA)-mediated therapies for neovascular AMD are under clinical investigation.^29^ Only one antisense oligonucleotide therapy has entered clinical trials for the treatment of AMD. The safety and pharmacodynamic profile of Sefaxersen (RO7434656), a GalNAc-conjugated 2’-MOE antisense oligonucleotide targeting hepatic complement factor B mRNA, has been investigated in a phase 1 study involving single or repeated subcutaneous administration of the drug, with a view to treat geographic atrophy secondary to AMD.^30^

Despite the complex aetiology of AMD, there is a significant overlap in pathological features between DHRD and AMD, including the drusen-associated accumulation of EFEMP1 in AMD patients.^9,31^ Therapeutic oligonucleotides developed for the treatment of AMD have not been investigated for the overlapping indications in DHRD. Currently, the clinical care of DHRD patients includes the use of low-vision aides to assist with daily living, and regular observation and retinal imaging to monitor drusen and detect complications such as choroidal neovascularisation (CNV). Interventions targeting complications that occur in some DHRD patients, including anti-VEGF injections (bevacizumab or ranibizumab) for CNV, photodynamic laser therapy and low-energy subthreshold nanosecond laser (2RT) treatment, have been trialled in a small number of DHRD patients.^32,33^ In comparison, our antisense oligonucleotide is the first to target the cause of DHRD at source and could potentially be used to treat all DHRD patients. Although DHRD is rare (c.1033C>T variant allele frequency of 1.23 × 10^−5^ in gnomAD database v4.1.0), it is a monogenic, single variant disorder with all DHRD patients worldwide harbouring the *EFEMP1* c.1033C>T, p.Arg(345Trp) variant. In an investigation of the spectrum of genetic variants in the most common genes causing inherited retinal disease in a large molecularly characterized cohort in the United Kingdom, the *EFEMP1* c.1033C>T, p.Arg(345Trp) variant was identified as one of the top twenty most prevalent variants identified.^34^ We conclude that our allele-specific antisense oligonucleotide could be a viable therapeutic option for DHRD patients, with the potential to alleviate the underlying pathology, including sub-RPE deposition of drusen-associated components and structural reorganisation of the RPE extracellular matrix, ultimately improving clinical outcomes.

## MATERIALS & METHODS

### EFEMP1 plasmids

*EFEMP1* (ENST00000355426.8) was amplified from cDNA prepared from retinal organoids as described in Sai *et al*. (2024).^35^ USER cloning technology (NEB, Hitchin, UK) was used according to the manufacturer’s instructions to insert the *EFEMP1* open reading frame excluding the stop codon into the mammalian expression vector p3xFLAG-CMV-14 (E7908) from Merck (Sigma-Aldrich, Gillingham, UK) and pmScarlet_C1 (Addgene, plasmid #85042), resulting in in-frame fusion of EFEMP1 with a C-terminal 3xFLAG tag (pEFEMP1-3xFLAG) and C-terminal mScarlet tag (pEFEMP1-mScarlet) respectively. The *EFEMP1* c.1033C>T, p.Arg(345Trp) variant and a Kozak sequence upstream of the *EFEMP1* open reading frame were introduced into pEFEMP1-mScarlet by site directed mutagenesis using the Q5 Site Directed Mutagenesis Kit (NEB) to generate pEFEMP1(R345W)-mScarlet. Primer sequences for *EFEMP1* amplification and sequencing are in Supplemental Table 5.

### ASO screening in HEK293T cells

HEK293 cells (ATCC) were cultured with DMEM (Life Technologies, Waltham, MA, USA) with 10% fetal bovine serum (Invitrogen, Thermo Fisher Scientific, Paisley, UK) at 37°C with 5% CO_2_. HEK293T cells were seeded in 24-well plates at a density of 0.7 × 10^5^ cells per well and transfected 24 hours later using Lipofectamine 2000 (Invitrogen, Thermo Fisher Scientific) as described by the manufacturer. HEK293T cells were transfected with both pEFEMP1(WT)-3xFLAG (800 ng) and pEFEMP1(R345W)-mScarlet (800 ng), and with control ASO (CTRL) (50 nM) or increasing concentrations (25 nM, 50 nM, 100 nM, 200 nM) of therapeutic ASO. Results were compared to untreated cells transfected with both plasmids (plasmids only). The RNA was extracted 48 hours post-treatment with the RNeasy Mini Kit (Qiagen, Manchester, UK) and cDNA synthesised from RNA samples using the Tetro cDNA Synthesis Kit (Bioline, Meridian Bioscience Europe, London, UK) according to the manufacturer’s protocol. Quantitative PCR (qPCR) was conducted using primer pairs specific for WT *EFEMP1,* R345W *EFEMP1* or total *EFEMP1* to quantify the levels of *EFEMP1* transcript using an Applied Biosystems QuantStudioTM 6 Flex real-time PCR system and the ΔΔCt method. Gene expression was normalized to the housekeeping genes *GAPDH* and *ACTB* and all values were expressed relative to the untreated ‘plasmid only’ condition. Primer sequences for *EFEMP1* qPCR are in Supplemental Table 5.

### Urine collection and renal epithelial cell isolation and expansion

Urine was collected from a DHRD patient harbouring the c.1033C>T, p.Arg(345Trp) variant in *EFEMP1*. Collection, isolation and expansion of renal epithelial cells was carried out similarly to protocols described by Zhou *et al.* (2012) and Hildebrand *et al.* (2016).^36,37^ Briefly, the urine sample (around 50 ml) was centrifuged at 400g for 10 minutes. The supernatant was discarded, and the pellet was washed with 10 ml of Dulbecco’s phosphate-buffered saline (Thermo Fisher Scientific) containing 1x Anti-Anti (Thermo Fisher Scientific) and centrifuged at 400g for 10 minutes. The pellet was resuspended in 2 ml of Primary Medium, consisting of DMEM/Ham’s F-12 nutrient mix (1:1) (Thermo Fisher Scientific), with 10% of fetal bovine serum (FBS; Thermo Fisher Scientific), Renal Cell Growth Medium (REGM) SingleQuot kit supplements (Lonza, Cambridge, UK) and 1x Anti-Anti. The cells were seeded into a single well of a 12-well plate coated with 0.1% gelatin. 1 ml of primary medium was added to the well at 24, 48 and 72 hours following plating, without removal of any media. Renal epithelial cell colonies appeared within 3 days. 96 hours post-seeding, most of the medium was aspirated and replaced by Proliferation Medium, consisting of a 1:1 mixture of Renal Epithelial Cell Growth Basal Medium (REBM) medium supplemented with REGM SingleQuots (Lonza) and DMEM high glucose (Thermo Fisher Scientific) supplemented with 10% FBS, 1% GlutaMAX, 1% non-essential amino acids (NEAA) and 1x Anti-Anti. Half media changes were performed every day. 14 days following urine collection, cells reached 90% density and were dissociated using TrypLE Express (Thermo Fisher Scientific) for 10 minutes at 37°C, before being seeded at a ratio of 1:4 onto gelatin coated wells. Cells were expanded for a maximum of four passages.

### Human iPSC reprogramming and culture

Renal epithelial cells passaged less than 3 times were used to generate iPSCs. 5 x 10^5^ cells were dissociated using TrypLE Express (Thermo Fisher Scientific) for 10 minutes at 37°C and electroporated with four integration-free episomal vectors: pCXLE-hOCT3/4-shp53-F (Addgene, plasmid #27077, pCXLEhSK (Addgene, plasmid #27078) containing SOX2 and KLF4, pCXLEhUL (Addgene, plasmid #27080) containing L-MYC and LIN28, and miRNA 302/367 plasmid (Gift from Dr J. A. Thomson, Regenerative Biology, Morgridge Institute for Research, Madison, Wisconsin, USA; Howden et al., 2015) using the AmaxaTM Basic NucleofectorTM Kit for primary mammalian epithelial cells, program T-020 (Lonza). Electroporated cells were seeded onto 1% Geltrex-coated (Thermo Fisher Scientific) plates and cultured in StemFlex media (Thermo Fisher Scientific). Media changes were carried out every other day. iPSC colonies were picked around day 14, expanded in Stem Flex media on Geltrex-coated 6-well plates and routinely passaged using Cell Dissociation Buffer (Thermo Fisher Scientific). iPSC line pluripotency was confirmed using iPSC-specific antibodies: anti-OCT4 (Abcam, Cambridge, UK), anti-Nanog (Abcam), anti SSEA4 (Cell Signalling Technologies, Cambridge, UK) and anti-TRA-1-80 (Invitrogen, Thermo Fisher Scientific) (Table S6).

### Generation of isogenic iPSC lines

To generate an isogenic control line (ISOCON), a 20nt gRNA (NGG PAM) and 127nt single stranded oligo deoxynucleotide (ssODN) template were designed for the correction of the *EFEMP1* c.1033C>T allele through CRISPR-mediated homology-directed repair (HDR). The ssODN repair template incorporated the desired correction and a synonymous PAM change to prevent further Cas9-mediated cleavage of the edited DNA. To generate an *EFEMP1* knock out (ISOKO) iPSC line, CRISPR-mediated non-homologous end joining (NHEJ) in the patient iPSC line was employed. A 20 nt gRNA with a canonical PAM (NGG) was designed to target exon 5 of *EFEMP1* (Table S2). The CRISPR RNA (crRNA) and ssODN (with phosphorothioate (PS) modifications to the ends) reagents were obtained from Integrated DNA Technologies (IDT, Bristol, UK). Sequences for crRNAs and ssODNs are in Supplementary Table 1 and 2 (Table S1 and S2). Trans-activating CRISPR RNA (tracrRNA), crRNA and ssDNA donors were resuspended in nuclease-free ddH_2_O to 100 µM. The crRNA and tracrRNA were combined and heated at 95°C for 5 minutes to form a 50 µM gRNA complex. 3 µl of gRNA complexes were then added to 2 µL of 61 µM Alt-R Cas9 enzyme (IDT) to generate ribonucleoprotein (RNP) complexes and incubated at room temperature (RT) for 20 minutes. iPSCs were cultured in Stemflex (Gibco, Thermo Fisher Scientific) with 10 uM ROCK inhibitor Y-27632 (STEMCell Technologies, Cambridge, UK) for 2 hours prior to single-cell dissociation with TrypLE (Gibco, Thermo Fisher Scientific). The dissociated cells (2 × 10^5^ cells per reaction) were centrifuged at 200g for 5 minutes and resuspended in P3 Primary cell nucleofector solution (Lonza) prior to the addition of 5 µl RNP solution, 2 µl of 100 µM ssODN (for CRISPR correction) and 4 µl Electroporation Enhancer. The iPSC-RNP mix was nucleofected using a P3 Primary Cell 4D-Nucleofector X Kit S (Lonza) using program CA-137. iPSC clones were expanded, and individual clones were mechanically isolated and plated into individual wells of a Geltrex-coated 12-well plate. Clonal iPSC lines were expanded and screened for the desired edit by Sanger sequencing (Source Bioscience).

### RPE differentiation

iPSCs were cultured to 90–100% confluency on Geltrex-coated plates (Thermo Fisher Scientific) and then directed to differentiate into retinal pigment epithelial (RPE) cells following the protocol described by Regent *et al.* (2019).^20^ The RPE differentiation medium comprised high-glucose Dulbecco’s modified Eagle’s medium (Thermo Fisher Scientific), supplemented with 50 μM β-mercaptoethanol (Thermo Fisher Scientific), 1× minimum essential medium–nonessential amino acids (Thermo Fisher Scientific), and knockout serum replacement (KSR, Thermo Fisher Scientific) at 20% from day 0 to day 50, which was reduced to 4% after passage 1. Additional supplements included 10 mM nicotinamide (Sigma-Aldrich) from days 0 to 7, 100 ng/ml Activin A (PeproTech, London, UK) from days 7 to 14, and 3 µM CHIR99021 (Cell Guidance Systems, Cambridge, UK) until passage 1. Media changes were carried out every 2 to 3 days. Around day 50, cells were treated with TrypLE Express Reagent (Thermo Fisher Scientific) at 37°C for 10 minutes. Visibly pigmented patches were manually isolated, and cells were further incubated for up to one hour with TrypLE Express to ensure dissociation. Finally, cells were seeded onto Geltrex-coated plates at a density of 100,000 cells per cm² for subsequent experiments.

### Antisense oligonucleotide gymnosis

Gymnotic ASO experiments took place by adding the desired ASO concentration into the media once a week. Half media changes were carried out 3 days later, with no ASO added.

### Immunostaining

iRPE monolayers grown in 12-well plates were washed with DPBS (Thermo Fisher Scientific) and were carefully removed using a cell scraper and transferred to bijou tubes where they were fixed in 4% paraformaldehyde (Thermo Fisher Scientific) for 10 mins at RT. The iRPE were washed 3x in DPBS and following the last wash, were incubated in 30% sucrose at 4°C overnight. iRPE monolayers were vertically embedded in OCT compound (CellPath, Newtown, UK) and frozen on dry ice. Blocks were stored at -80°C prior to sectioning at 14 μm onto charged slides using a Leica CM1850 cryostat. Cells grown on chamber slides were washed with PBS (Thermo Fisher Scientific) and fixed in 4% paraformaldehyde (Thermo Fisher Scientific) for 10 mins at RT. Cells grown in chamber slides and vertically-embedded iRPE were permeabilised using 0.1% TritonX100 (Sigma-Aldrich) for 15 minutes before being blocked using a solution containing 0.3% bovine serum albumin (BSA, Sigma-Aldrich), 10% FBS (Thermo Fisher Scientific) for 1 hour at RT. Primary antibodies (Table S6) were diluted at the appropriate concentration in blocking solution and incubated for 1 hour at RT. Samples were washed 3x with PBS (Thermo Fisher Scientific) and subsequently incubated with secondary antibodies (Table S7) that reacted against the primary antibody species for 1 hour at RT, before being washed 3x with PBS. For BODIPY 493/503 (Thermo Fisher Scientific) staining, samples were incubated for 15 minutes using a 10 mM BODIPY stock that was diluted 1:1000 following 2x washes with PBS. All samples were incubated with 4,6-diamidino-2-phenylindole (DAPI; 2 mg/mL) (Invitrogen, Thermo Fisher Scientific) in PBS for 5 minutes, then washed 3x with PBS. Slides were mounted in fluorescence mounting media (Dako, Agilent, CA, USA). All images were acquired using a Zeiss LSM700 Confocal Microscope and prepared using ImageJ, Adobe Photoshop and Adobe Illustrator.

### Western blotting

iRPE were washed with PBS and lysed on ice for 30 minutes in 80-120μl cold RIPA buffer (50mM Tris HCL, pH 7.5; 150mM NaCl; 1mM EDTA; 1% NP-40; 0.5% sodium deoxycholate; 0,1% SDS) supplemented with 2% protease inhibitor cocktail (Sigma-Aldrich). The samples were briefly sonicated before centrifuging at 4°C for 1 minute at 13k RPM. Protein concentration was quantified using Pierce BCA Protein Assay kit (Thermo Fisher Scientific) as per manufacturer’s instructions. Protein samples (20 µg) were mixed with 4x sample buffer before being heated at 100°C for 3 minutes to facilitate protein denaturation. Samples were resolved on precast 4–20% gradient gels (Bio-Rad, Watford, UK) at 100 V until the dye reached the end of the gel. Pageruler Plus protein ladder (Thermo Fisher Scientific) was used to provide a reference for protein size. Proteins were transferred to a nitrocellulose membrane by wet transfer for 90 minutes, using standard protocols. Membranes were blocked overnight at 4°C in 5% skimmed milk powder in PBS + 0.1%Tween-20 (PBS-T), then incubated for 1 hour in primary antibody, before undergoing 3x 10-minute washes with PBS-T. Membranes were then incubated in species-specific HRP-conjugated secondary antibody for 1 hour before undergoing 3x 10-minute washes with PBS-T. All antibodies are listed in supplemental table 6 and 7 (Table S6). Membranes were developed using Clarity MAX Western ECL (Bio-Rad) for 5 min. Membranes were imaged using a ChemiDoc MP Imaging System (Bio-Rad). Images were formatted using Image Lab (Bio-Rad). Three independent protein isolates were used per condition.

### Transmission electron microscopy

iRPE cells cultured on transwell filters were fixed in EM fixative (2% gluteraldehyde, 2% paraformaldehyde in 0.1M sodium cacodylate buffer, pH 7.4) for 30 min at room temperature and post-fixed with 1% osmium tetroxide/1.5% potassium ferricyanide in 0.1 M sodium cacodylate buffer for 1 hour at 4°C. Cells were stained with 2% uranyl acetate replacement solution (UA Zero) for 1 hour, dehydrated through an ethanol series, and infiltrated with TAAB-812 epoxy resin (TAAB Laboratories, Aldermaston, UK) prior to polymerization overnight at 60°C. Ultrathin sections were imaged on a JEOL 1400Plus EM (JEOL ltd, Tokyo, Japan) fitted with an Advanced Microscopy Technologies (AMT) NanoSprint12 camera (AMT Imaging Direct, Woburn, MA, USA). Quantitation of accumulations between the basal membrane and underlying transwell membrane was conducted manually and blind to the condition. ISOCON iRPE, 40 cells (∼10 μm/cell); R345W iRPE, 25 cells (∼ 7.8 μm/cell); ASO1.1-treated R345W iRPE, 22 cells (∼9.7 μm/cell).

### Processing of DNA and RNA from iRPE

Total gDNA from cell samples was extracted using the Wizard gDNA Purification Kit (Promega) following the manufacturer’s guidelines. Total RNA from cell samples was extracted using the RNeasy Micro Kit (Qiagen) following the manufacturer’s guidelines. cDNA was synthesised (Tetro cDNA synthesis, Bioline) using equal quantities of total RNA for each sample (1 μg) and carried out according to the manufacturer’s guidelines.

### Quantitative PCR (qPCR) of iRPE samples

qPCR was used to determine relative levels of *EFEMP1* gene expression (normalised to the housekeeping genes *ACTB* and *GAPDH*), by measuring the fluorescence emitted by DNA bound to the intercalating dye SYBR green. qPCR reactions were assembled using 2x LabTaq Green Hi Rox Master Mix (Labtech, Heathfield, UK) and conducted with an Applied Biosystems QuantStudio ™ 6 Flex real-time PCR system.

### Next generation sequencing

Universal tagged primers for MiSeq (Illumina) high-throughput sequencing (HTS) are listed in Supplemental Table 5 (Table S5). PCR amplification was carried out using High-Fidelity 2x master mix (NEB). Products were gel-extracted (Monarch® DNA Gel Extraction Kit, NEB) and 300-500ng of product used for subsequent steps. Processing of the samples using MiSeq Reagent Nano Kit v2 was carried out at the UCL Cancer Institute CAGE Facility.

## Data visualisation and statistical analysis

Statistical analyses were carried out using RStudio (R v4.2.1). One way ANOVAs were carried out using the base R aov function and post-hoc analyses using the RStudio package rstatix (v0.7.2). All graphs presented were generated using the RStudio package using the RStudio package ggplot2 (v3.4.1).

## DATA AVAILABILITY

The data related to this study are fully documented in the paper or in the Supplemental materials.

## ACKNOWLEDGEMENTS

The authors gratefully acknowledge funding from the Macular Society (20-RG-04 to J.v.d.S (PI), A.-J.F.C. (Co-I), M.E.C. (Co-I) and M.M. (Co-I)). This research was supported by the National Institute for Health and Care Research Biomedical Research Centre at Moorfields Eye Hospital and UCL Institute of Ophthalmology and the Wellcome Trust (099173/Z/12/Z to M.M. and 205041/Z/16/Z to M.E.C.). pmScarlet_C1 was a gift from Dorus Gadella (Addgene plasmid # 85042). miRNA 302/367 plasmid was a gift from Dr J. A. Thomson, Regenerative Biology, Morgridge Institute for Research, Madison, Wisconsin, USA (Howden et al., 2015). pCXLE-hOCT3/4-shp53-F (Addgene, plasmid #27077), pCXLEhSK (Addgene, plasmid #27078) and pCXLEhUL (Addgene, plasmid #27080) were a gift from Shinya Yamanaka. We thank the Cancer Genomics Engineering (CAGE) Facility, UCL Cancer Institute, for the MiSeq preparation and sequencing. The CAGE Facility is supported by the BRC, Welton Foundation and in part by the Cancer Research UK – UCL Centre.

## AUTHOR CONTRIBUTIONS

Conceptualization, J.v.d.S.; Data curation, F.O.R.; E.R.E.; J.v.d.S.; Formal analysis, F.O.R.; B.S-P.; E.R.E., J.v.d.S.; Funding Acquisition, J.v.d.S.; A-J.F.F.; M.E.C.; M.M. Investigation, F.O.R.; B.S-P.; E.R.E.; Methodology, F.O.R.; B.S-P.; E.R.E.; J.v.d.S.; Project Administration, J.v.d.S.; Resources, N.A.; A.R.W.; A.-J.F.C; M.E.C.; M.M.; Supervision, J.v.d.S; Software, F.O.R.; Validation, F.O.R.; B.S-P.; E.R.E., J.v.d.S; Visualization; F.O.R.; B.S-P.; E.R.E., J.v.d.S.; Writing – original draft; F.O.R.; J.v.d.S.; Writing – review and editing, F.O.R.; B.S-P.; E.R.E., N.A.; A.R.W.; A.-J.F.C; M.E.C.; M.M.; J.v.d.S.

