## Supplemental Data for "Antisense oligonucleotide allele-specific targeting of EFEMP1 in a patient-derived model of Doyne honeycomb retinal dystrophy"

**Supplemental Table 1: CRISPR Correction of c.1033C>T: crRNA and ssODN sequences**

| Desired change | crRNA (5'-3') | GC% | ssODN (5'-3') |
|---|---|---|---|
| c.1033T>C | GACCACAAATGAATGCTGGG | 50% | TTCTCTGGTGTTAGAAT<br>GTAGGGATCTTGACAA<br>GGATTTCGTGGATAACA<br>ACGGAAGCCGCCATGA<br>TAATTCCAACACATTTC<br>ATC**T**TCCCGGCATTCAT<br>TAGTGGTCTCACACTCA<br>TTTATATCTG |

**cRNA**: Target residue (T): large font size. **ssODN**: Complementary to cRNA (underlined). Target repair (G): large font size. Synonymous PAM change (T): bold

**Supplemental Table 2: CRISPR *EFEMP1* Knockout: cRNA Sequence**

| Resulting change | cRNA (5'-3') | GC% |
|---|---|---|
| c.215_216insG,<br>p.Lys73GlufsX81 | TCCGAAAACAGCCCAGATTA | 45% |

**Supplemental Table 3: Cas-OFFinder Analysis of *In Silico* Predicted Off Targets**

| Guide purpose | Site of potential off-target editing | Sequence of potential binding site | Mismatches |
|---|---|---|---|
| CRISPR Correction of c.1033C>T | chr6:51,829,169-51,829,191 | crRNA:<br>GACCACAAATGAATGCTGGG<br>DNA:<br>tAgCACAAATcAATGCTGGG-TGG | 3 |
| CRISPR Correction of c.1033C>T | chr16:64,364,348-64,364,370 | crRNA:<br>GACCACAAATGAATGCTGGG<br>DNA:<br>GAtCACAAgTGcATGCTGGG-AGG | 3 |
| CRISPR Correction of c.1033C>T | chr8:144,720,814-144,720,836 | crRNA:<br>GACCACAAATGAATGCTGGG<br>DNA:<br>GAgCACAAAgGAATtCTGGG-TGG | 3 |
| CRISPR Correction of c.1033C>T | chr10:29,247,464-29,247,486 | crRNA:<br>GACCACAAATGAATGCTGGG<br>DNA:<br>GACaAgAAATGtATGCTGGG-AGG | 3 |
| CRISPR Correction of c.1033C>T | chr1:211,488,887-211,488,909 | crRNA:<br>GACCACAAATGAATGCTGGG | 3 |

| | | | |
|---|---|---|---|
| | | DNA: GACCtCAgATGcATGCTGGG-TGG | |
| CRISPR Correction of c.1033C>T | chr5:155,412,417-155,412,439 | crRNA: GACCACAAATGAATGCTGGG DNA: agaCACAAAaGAATGCTGGG-CGG | 4 |
| CRISPR Correction of c.1033C>T | chr3:8,212,410-8,212,432 | crRNA: GACCACAAATGAATGCTGGG DNA: ttCaACAcATGAATGCTGGG-GGG | 4 |
| CRISPR Correction of c.1033C>T | chr10:73,553,379-73,553,401 | crRNA: GACCACAAATGAATGCTGGG DNA: ctCCACAgATGcATGCTGGG-CGG | 4 |
| CRISPR Correction of c.1033C>T | chr6:10,342,654-10,342,676 | crRNA: GACCACAAATGAATGCTGGG DNA: ttCCACAcATGAATtCTGGG-GGG | 4 |
| CRISPR Correction of c.1033C>T | chr14:58,766,075-58,766,097 | crRNA: GACCACAAATGAATGCTGGG DNA: cAaggCAAATGAATGCTGGG-AGG | 4 |
| *EFEMP1* knockout | chr4:72,743,031-72,743,053 | crRNA: TCCGAAAACAGCCCAGATTA DNA: TCaGAAAcCAGCtCAGATTA-TGG | 3 |
| *EFEMP1* knockout | chr6:135,177,560-135,177,582 | crRNA: TCCGAAAACAGCCCAGATTA DNA: ggtGAAAACAGaCCAGATTA-GGG | 4 |
| *EFEMP1* knockout | chr4:93,297,559-93,297,581 | crRNA: TCCGAAAACAGCCCAGATTA DNA: atCtAAAAaAGCCCAGATTA-GGG | 4 |
| *EFEMP1* knockout | chr19:17,948,974-17,948,996 | crRNA: TCCGAAAACAGCCCAGATTA DNA: aaCGAcAAtAGCCCAGATTA-GGG | 4 |
| *EFEMP1* knockout | chr14:76,683,368-76,683,390 | crRNA: TCCGAAAACAGCCCAGATTA DNA: gCtaAAtACAGCCCAGATTA-TGG | 4 |

| | | | |
|---|---|---|---|
| *EFEMP1* knockout | chr3:170,532,773-170,532,795 | crRNA: TCCGAAAACAGCCCAGATTA<br>DNA: aCaGAAAAaAGCCCtGATTA-AGG | 4 |
| *EFEMP1* knockout | chr8:134,919,582-134,919,604 | crRNA: TCCGAAAACAGCCCAGATTA<br>DNA: aCCaAgAACAtCCCAGATTA-GGG | 4 |
| *EFEMP1* knockout | chr1:52,983,771-52,983,793 | crRNA: TCCGAAAACAGCCCAGATTA<br>DNA: aCCcAAgAaAGCCCAGATTA-TGG | 4 |
| *EFEMP1* knockout | chr9:119,373,155-119,373,177 | crRNA: TCCGAAAACAGCCCAGATTA<br>DNA: aCCcAAAgCAcCCCAGATTA-AGG | 4 |
| *EFEMP1* knockout | chr2:163,975,413-163,975,435 | crRNA: TCCGAAAACAGCCCAGATTA<br>DNA: TtCtAAAACAtgCCAGATTA-AGG | 4 |

**Supplemental Table 4: ASO Sequences**

| Name | Sequence (5'-3') | Chemistry |
|---|---|---|
| CTRL ASO | CACCCCCATTCTTCAGCC | PS backbone; 2'-O-MOE |
| ASO1 | UCAUCCTCCCAGCAUUCA | PS backbone; 2'-O-MOE; LNA* |
| ASO2 | UCAUCCTCCCAGCAUUCA | PS backbone; 2'-O-MOE, LNA** |
| ASO3 | AUCCUCCCAGCATUCAUU | PS backbone; 2'-O-MOE; LNA* |
| ASO4 | AUCCUCCCAGCATUCAUU | PS backbone; 2'-O-MOE; LNA** |
| ASO1.1 | CATCCTCCCAGCATTC | PS backbone; 2'-O-MOE; LNA* |
| ASO1.2 | TCATCCTCCCAGCAT | |
| ASO1.3 | ATCCTCCCAGCATTC | |
| ASO1.4 | ATCCTCCCAGCATT | |

*PS, phosphorothioate; 2'-O-MOE, 2'-O-methoxyethyl; LNA*, single linked nucleic acid; LNA**, two linked nucleic acids.*

**Supplemental Table 5: Primers**

| **EFEMP1 PCR Primers** | |
|---|---|
| **Name** | **Sequence (5'-3')** |
| EFEMP1-201_1F (Forward) | TGTGCTGTGCAAGGAACTCT |

| | |
|---|---|
| EFEMP1-201_2F (Forward) | GCTGTGCAAGGAACTCTGCT |
| EFEMP1-201_1R (Reverse) | TTGGCTGACTTAAATGCCTGT |
| **Sequencing Primers** | |
| EFEMP1 c.1033C>T cDNA F (Forward) | TGCAGAACCTCAAGCTACCTGTGTC |
| EFEMP1 c.1033C>T cDNA R (Reverse) | GGGCAAACACATCGGTTCTCTGG |

| **qPCR Primers** | | |
|---|---|---|
| **Name** | **Forward (5'-3')** | **Reverse (5'-3')** |
| MScarlet | GAGTTCATGCGGTTCAAGGT | ACATGAACTGAGGGGACAGG |
| 3xFLAG | GTATAGGGACCTTCCGCACA | GGAGGGGTCACAGGGATG |
| EFEMP1 | GTTTCCTGCTGAGGCTGTTC | CAGGACACCGAAGAAACCAT |
| GAPDH | CCCCACCACACTGAATCTCC | GGTACTTTATTGATGGTACATGACAAG |
| ACTB | CCAACCGCGAGAAGATGA | CCAGAGGCGTACAGGGATAG |

| **High Throughput Sequencing (HTS) Primers** | | | |
|---|---|---|---|
| **Forward MiSeq primer (5'-3')** | **Reverse MiSeq primer (5'-3')** | **Amplicon Size** | |
| | | **bp w/o tags** | **bp w/ tags** |
| **TCGTCGGCAGCGTCAGATGTGTATAAGAGACAG** TGCAGAACCTCAAGCTACCTGTGTC | **GTCTCGTGGGCTCGGAGATGTGTATAAGAGACAG** GGGCAAACACATCGGTTCTCTGG | 248 | 315 |
| **TCGTCGGCAGCGTCAGATGTGTATAAGAGACAG** CCTTCCTTGCAAACAGAATCTGCC | **GTCTCGTGGGCTCGGAGATGTGTATAAGAGACAG** GCAGTTTGGCTTGGTAAGACCAG | 276 | 343 |

**Supplemental Table 6: Primary Antibodies**

| **Antigen** | **Host** | **Supplier** | **Catalogue number** | **Dilution** |
|---|---|---|---|---|
| ZO-1 | Rabbit | Thermo Fisher Scientific | 61-7300 | 1:1000 (WB)<br>1:200 (IF) |
| PMEL | Mouse | Agilent | M0634 | 1:50 (IF) |
| MERTK | Rabbit | Abcam | ab52968 | 1:500 (WB)<br>1:200 (IF) |
| EZRIN | Mouse | Thermo Fisher Scientific | MA5-13862 | 1:500 (WB)<br>1:200 (IF) |
| GAPDH | Mouse | Proteintech | 60004-1-Ig | 1:10,000 (WB) |
| Collagen IV | Goat | Bio-techne | NBP1-26549 | 1:200 (IF) |

| | | | | |
|---|---|---|---|---|
| APOE | Mouse | Bio-techne | NB110-60531 | 1:200 (IF) |
| Fibulin 3 | Mouse | Santa Cruz Biotechnology | sc-33722 | 1:250 (IF) |
| OCT4 | Rabbit | Abcam | Ab19857 | 1:1000 (IF) |
| NANOG | Mouse | Invitrogen | MA1-017 | 1:200 (IF) |
| SSEA4 | Mouse | Cell Signalling Technologies | 4755 | 1:500 (IF) |
| TRA181 | Mouse | Cell Signalling Technologies | 4745 | 1:1000 (IF) |

**Supplemental table 7: Secondary Antibodies**

| **Antigen** | **Host** | **Fluorophore** | **Supplier** | **Catalogue number** | **Dilution** |
|---|---|---|---|---|---|
| Rabbit IgG | Donkey | 488 | Invitrogen | A32790 | 1:1000 |
| Mouse IgG | Donkey | 555 | Invitrogen | A31571 | 1:1000 |
| Goat IgG | Donkey | 488 | Invitrogen | A-21432 | 1:1000 |
| Rabbit-HRP | Goat | - | Invitrogen | 31460 | 1:30,000 |
| Mouse-HRP | Goat | - | Invitrogen | 31430 | 1:30,000 |

SUPPLEMENTAL FIGURES

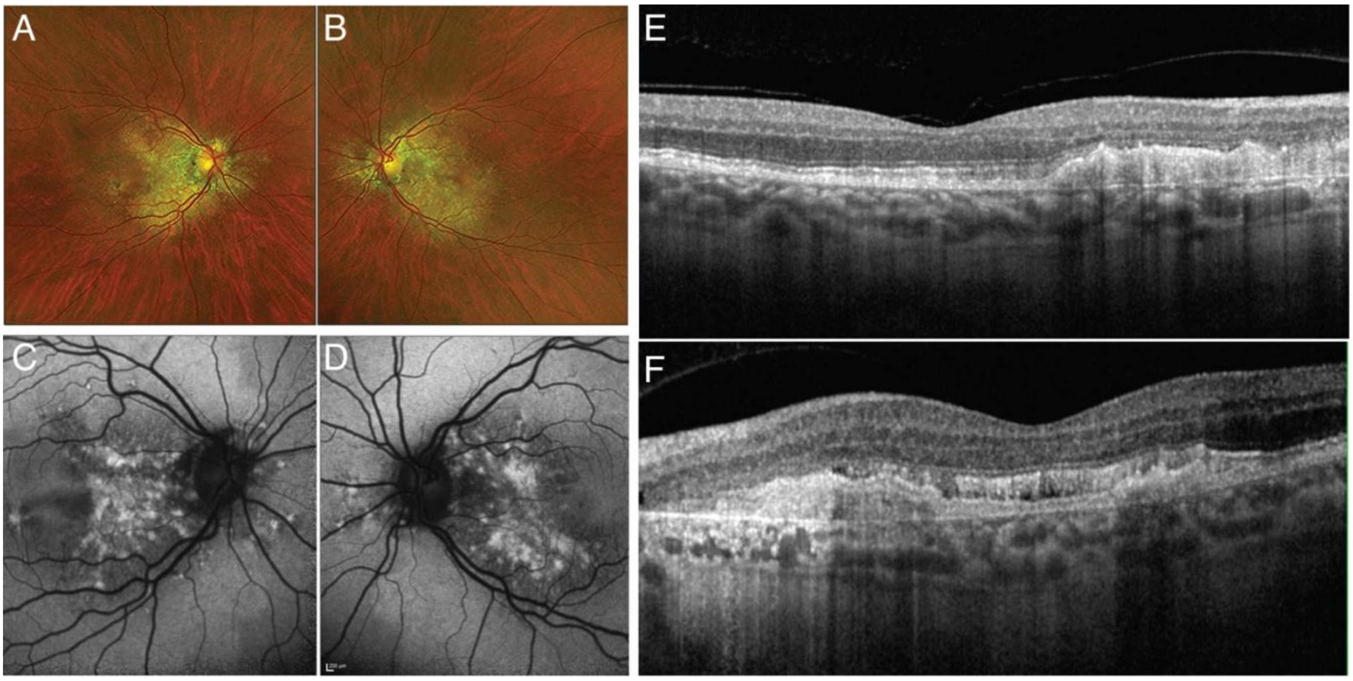


**Supplemental Figure 1: Clinical features of DHRD patient at 60 years of age.**
Related to Figure 1.
(A) Ultra-widefield fundus pseudocolour image (Optos scanning laser ophthalmoscope) of right eye.
(B) Ultra-widefield fundus pseudocolour image (Optos scanning laser ophthalmoscope) of left eye.
(C) Autofluorescence image of right eye (HEYEX blue light AF).
(D) Autofluorescence image of left eye (HEYEX blue light AF).
(E) Heidelberg Spectralis OCT image of right eye.
(F) Heidelberg Spectralis OCT image of left eye.

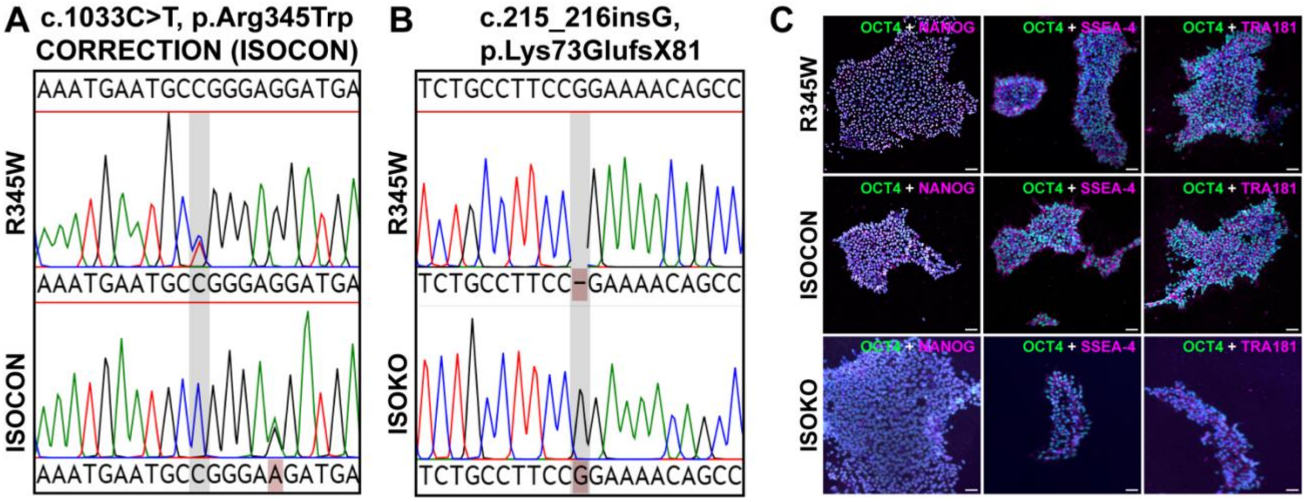


**Supplemental Figure 2: CRISPR-Cas9 genome editing of EFEMP1 isogenic control (ISOCON) and knockout (ISOKO) iPSC**

Related to Figure 1.

(A) Sanger sequencing chromatogram of gDNA from patient-derived R345W iPSC (top) and ISOCON iPSC (bottom) following CRISPR-Cas9 HDR to correct *EFEMP1* c.1033C>T (grey bar). The synonymous change in the PAM (AGG-AAG) is coloured pink.

(B) Sanger sequencing chromatogram of gDNA from patient-derived R345W iPSC (top) and ISOKO iPSC (bottom) following CRISPR-Cas9 NHEJ to knockout the *EFEMP1* gene. The grey bar highlights the insertion of a single homozygous ‘G’ at c.125.

(C) Characterization of ISOCON and IOSKO iPSC clones. Immunocytochemistry analysis of pluripotency markers Oct4, Tra-1-81, Nanog and SSEA-4 in patient-derived R345W, ISOCON and ISOKO iPSC. Scale bars, 10 μm.
